# Quantification of collagen and associated features from H&E-stained whole slide pathology images across cancer types using a physics-based deep learning model

**DOI:** 10.1101/2025.03.17.643273

**Authors:** Tan H. Nguyen, Jun Zhang, Jennifer Hipp, Guillaume Chhor, Michael Griffin, Nhat Le, Deeksha Kartik, Yibo Zhang, Mohammad Mirzadeh, Julia Varao, Jim Allay, Morgan Sweeney, Veronica Rivera, Brea Johnson, Jacqueline Brosnan-Cashman, Matthew Bronnimann, Harsha Pokkalla, Ben Glass, Andrew H. Beck, Justin Lee, Robert Egger

## Abstract

**Background:** Collagen is the major component of the extracellular matrix (ECM). Collagen structural organization undergoes significant transformation during tumorigenesis. The visualization of collagen in histological tissue sections would aid in the study of tumor growth, encapsulation, and invasion. However, such visualization requires the use of special stains such as Picrosirius Red (PSR) or Masson’s Trichrome (MT), or more recently, second-harmonic generation imaging (SHG) in unstained tissue sections. However, PSR and MT both suffer from significant inter- (and intra-) lab stain variabilities, and SHG, while considered a ground truth by many, suffers from issues of system complexity/reliability, cost, and speed/throughput. These technical hurdles limit more widespread assessment of collagen in tissue samples.

**Methods:** Using high-contrast, high-throughput polarization imaging on PSR-stained slides to generate ground truth training polarization images, we developed a deep learning model (iQMAI) to infer the presence of collagen directly from hematoxylin and eosin (H&E)-stained whole-slide images (WSIs) with high specificity. After iQMAI inference across WSIs, individual collagen fibers were extracted, and features describing overall collagen intensity and fiber morphology were computed. iQMAI pixel-and feature-wise outputs were compared to ground truth polarization imaging to assess model performance. The trained iQMAI model was deployed on H&E-stained WSI from the TCGA LUAD, LUSC, LIHC, and PAAD datasets for evaluation. iQMAI-derived collagen features were compared to tissue composition, gene expression, and overall survival.

**Results:** The iQMAI model shows significant generalization across multiple indications. iQMAI collagen predictions were similar to polarization imaging measurements of the same sample, with a mean structural similarity index (SSIM) of 0.84 (95% CI 0.69-0.93), a mean patch-wise RMSE of 0.04 (95% CI 0.02-0.08), and a linear correlation (R^2^=0.93). Comparing features of the collagen fibers extracted from iQMAI vs. polarization images yielded similar linear correlations between computed fiber tortuosity, length, width, and relative angle. The relationship between collagen fiber density and fibroblast density was distinct in non-small cell lung cancer (LUAD and LUSC), hepatocellular carcinoma (LIHC), and pancreatic ductal adenocarcinoma (PAAD). In PAAD, fiber density and fiber width were both negatively associated with the LRRC-15 gene expression signature, and increased fiber width was associated with longer overall survival.

**Conclusions:** iQMAI is a deep learning model that accurately predicts collagen from an H&E-stained WSI, allowing for spatially resolved quantification of collagen morphology and enabling investigation of the interplay between collagen and other TME components. We demonstrate an example of the utility of iQMAI-based collagen assessment in PAAD, where collagen features are associated with immunosuppressive cancer-associated fibroblasts and overall survival. Understanding the relationship between collagen, the tumor microenvironment composition, and disease progression may aid the development of effective immunotherapies in PAAD and other cancer types.

## Introduction

A tumor develops as a complex ecosystem comprising neoplastic epithelial cells and a heterogeneous tumor microenvironment (TME). Many varied cell types, such as cancer-associated fibroblasts (CAFs) and immune cells, are present within the TME, as is a supporting extracellular matrix (ECM).

In many cancers, a desmoplastic stroma forms, characterized by excessive deposition of ECM that is similar to chronic fibrosis when assessed by histological or molecular analysis.^1^ As such, the most abundant component of the cancer ECM is fibrillar collagen, which is produced by activated CAFs.^2^ Notably, the ECM can be altered in a variety of ways during cancer progression, including increased ECM production or degradation, changes in biochemical properties of intermolecular interactions, and altered matrix organization,^1^ leading to an aligned and anisotropic organization of collagen in many tumors.^3,4^

Collagen dynamics in the tumor ECM have been shown to impact tumor behavior in many ways, ultimately affecting patient outcome.^1^ In particular, the organization of collagen within the matrix is garnering increasing attention. The structure of the matrix, including fiber stiffness, impacts a cancer cell’s ability to invade through the matrix during metastasis.^5^ Indeed, in breast cancer, stromal collagen aligned perpendicular to the cancer epithelium is associated with reduced survival and increased metastasis.^6–8^ Aligned collagen fibers are also associated with poor survival in patients with head and neck squamous cell carcinoma and pancreatic ductal adenocarcinoma,^9,10^ while collagen width and tortuosity contribute to prognosis in lung cancer.^11^

Stromal collagen organization also impacts anti-tumor immunity. *In vitro* studies have demonstrated that high-density collagen slows the migration of T cells, which utilize aligned collagen fibers for migration.^12–16^ Furthermore, high collagen levels in lung cancer have been shown to be associated with poor response to checkpoint inhibition, while a cancer-associated ECM signature was elevated in non-responders to PD-1 inhibition in melanoma and bladder cancer.^17,18^

While collagen is widely understood to be an important component of cancer development and progression, the detection and visualization of collagen in clinical histological tissue specimens has been a challenge. Collagen can be imaged in tissue sections using brightfield microscopy with histological stains such as Movat’s pentachrome,^19^ Masson’s trichrome,^20^ or polarization-based microscopy,^21–23^ potentially incorporating Picrosirius red staining.^24–26^ Currently, second harmonic generation (SHG)^27–29^ is considered the gold standard for collagen imaging due to its ability to highlight non-centrosymmetric structures of the collagen molecules, as well as its high specificity and impressive depth sectioning. However, SHG is incompatible with clinical laboratory usage due to the need for specialized equipment and most importantly, its requirement of an unstained slide, similar to other birefringence-sensitive approaches such as LC-PolScope,^30^ polychromatic polarization imaging,^31^ and Mueller matrix microscopy.^32,33^

Advances in machine learning (ML) have great potential to aid the detection of collagen in tissue. ML approaches have been used for a myriad of purposes in cancer pathology, including classifying polarization-based images into normal vs. tumor,^34^ tumor diagnosis in multiple organs,^35–37^ and prediction of cancer recurrence after treatment.^33,38,39^ Additional applications of such algorithms have been applied to quantitation of fibrosis, such as quantifying scar tissue after surgery,^40^ staging liver fibrosis,^41^ and evaluating the mechanical properties of tissue.^42,43^ Furthermore, computer vision algorithms have been developed to allow morphological features of collagen fibers to be extracted and computed in images derived from SHG or collagen stains.^44^ However, this algorithm is not scalable for use on whole-slide images (WSI) and is incompatible with H&E-stained images, limiting its potential clinical utility. Finally, these approaches do not allow a spatial understanding of collagen in the TME, as they require a separate slide from the standard hematoxylin and eosin (H&E) staining and obscure concurrent spatial analyses of TME components (e.g., cell types, tissue regions).

Here, we introduce a novel technique for the detection of collagen in histological specimens: a machine learning-based approach to predict collagen structures on digital WSI of H&E-stained slides at scale. This model was trained using automated ground truth labels of collagen signals generated using polarized light imaging aligned with H&E-stained WSI from 4.3 million images derived from 15 indications. Importantly, features describing collagen levels and collagen fiber morphology can be computed from these techniques and used to examine clinically-relevant biological questions about the role of collagen in the TME in a spatially-resolved manner.

## Results

### Automated generation of collagen ground truth data for model training

Given the specialized imaging and stains traditionally required for the detection of collagen in histologic specimens, we aimed to generate an ML model to predict the presence of collagen in an H&E-stained WSI. Traditionally, ML models to identify and segment substances in histological images have necessitated manual substance annotations for model training. However, the resolution necessary to accurately identify collagen fibers renders manual annotations of these structures untenable, particularly from an H&E image. When considering other sources of ground truth collagen fibers, we considered other microscopy techniques. However, due to the use of serial sections and a low imaging throughput, SHG does not allow pixel-level registration to an H&E image and is not scalable to a WSI. Meanwhile off-the-shelf polarization microscopy techniques are fiber orientation-dependent, have a suboptimal signal to noise ratio, and identify false positive tissue substances (e.g., blood vessels) as harboring collagen signal. Therefore, to generate comprehensive model training and evaluation data describing the collagen content in H&E WSI at pixel-level resolution, we developed a custom imaging technique based on the principle of polarization microscopy, termed quantitative multimodal anisotropy imaging (QMAI).

Briefly, H&E-stained histologic slides were imaged with commercial brightfield WSI scanners, then restained with PSR and imaged using a microscope modified to accommodate multispectral, brightfield, and polarization imaging to visualize tissue components with birefringence (**Figure 1A**). The combination of PSR and polarization imaging ensures high sensitivity and specificity, as previously demonstrated^26,45^. This process allowed the polarization image, depicting collagen signal, to be registered to the H&E-stained image using the multispectral image. We directly compared the outputs of this technique to those of SHG in a neighboring unstained tissue section. While minor discrepancies in tissue architecture were observed, likely reflecting the inherent differences between neighboring tissue sections, we observed a high degree of similarity between the outputs of SHG and polarization imaging using PSR-stained slides, highlighted in **Figure 1B**.

**Figure 1.**
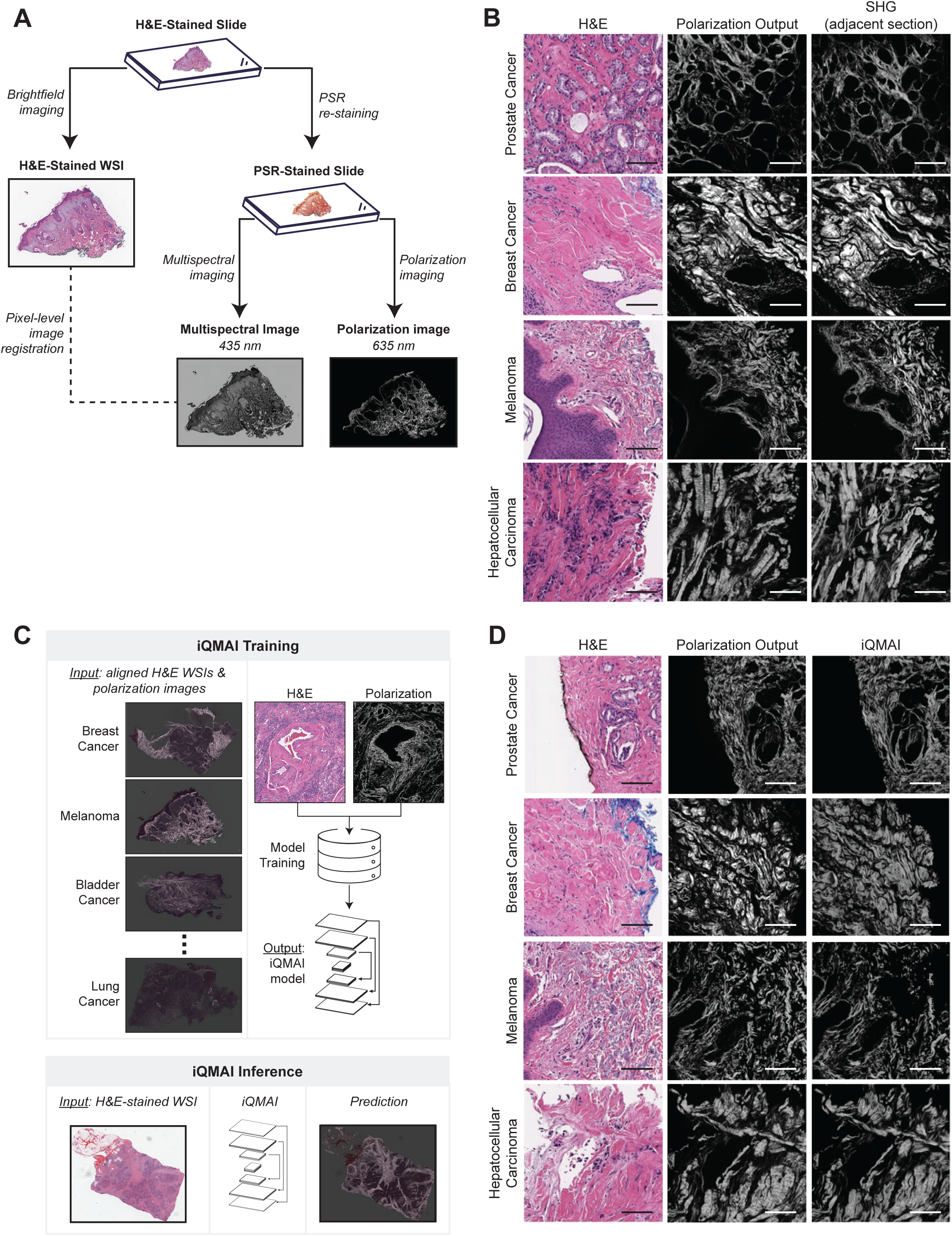
Training of a model to predict collagen signal in histopathology slides using images derived from polarization-based microscopy. A) Workflow to generate images to be used for model training. B) Comparison of polarization images for ground truth generation to SHG images from neighboring sections. Scale bars indicate 100 μm. C) Workflow for iQMAI training and inference. D) Comparison of iQMAI predictions to polarization imaging of the same slide. Scale bars indicate 100 μm.

### Collagen detection model development and evaluation

Having determined that our polarization imaging approach highlights collagen structures in a comparable manner to SHG, we proceeded to use images obtained through this modality to train a model to predict collagen from H&E-stained WSI. A dataset consisting of H&E-stained slides derived from fifteen cancer indications (N=337) was split into training (N=244; 72%), validation (N=47; 14%), and held-out test cohorts (N=46; 14%). Slides were used to generate ground truth images, as follows: H&E-stained slides were scanned to generate WSI then were re-stained with PSR to improve specificity for polarization microscopy. Resulting polarization WSI were registered to the H&E brightfield images, and the measured collagen intensity signal was used as an exhaustive “super-annotation” map of the WSI. These super-annotations were then used to train the model to infer the polarization imaging collagen signal from H&E-stained WSI (**Figure 1C**). After an examination of models generated with varying training set sizes (**Supplementary Table 1**), reconstruction losses (**Supplementary Table 2**), and weight combinations (**Supplementary Table 3**), the model trained with 0.01*ɭ*_LPIPS_ + 0.99*ɭ*_DICE_ was deemed to show optimal performance and was used for subsequent studies herein (**Supplementary Figure 1**). Henceforth, we refer to the model as inferred quantitative multimodal anisotropy imaging (iQMAI).

A qualitative assessment of iQMAI performance was performed by comparing the iQMAI predictions to polarization and SHG outputs on the same slides assessed previously. This comparison revealed that iQMAI predictions closely matched collagen derived from physical imaging (**Figure 1D**).

We sought to further evaluate the model in a quantitative manner. To do so, we compared the collagen outputs from iQMAI to those from the ground-truth polarization images. For a comparison of collagen intensity at the patch level, image frames (300 μm x 300 μm; N=2293) were randomly sampled from the usable tissue across 46 slides. For each frame, iQMAI and ground truth patches were extracted and compared. A comparison of the average patch-wise pixel intensities between the two outputs revealed a mean structure similarity index^46^ of 0.84 (95% CI 0.69-0.93), a mean patch-wise root-mean-square error of 0.04 (95% CI 0.02-0.08), and a linear relationship (R^2^=0.93; **Figure 2A**). Using these patch-wise average intensities, the ICC(2,1) was 0.96.

**Figure 2.**
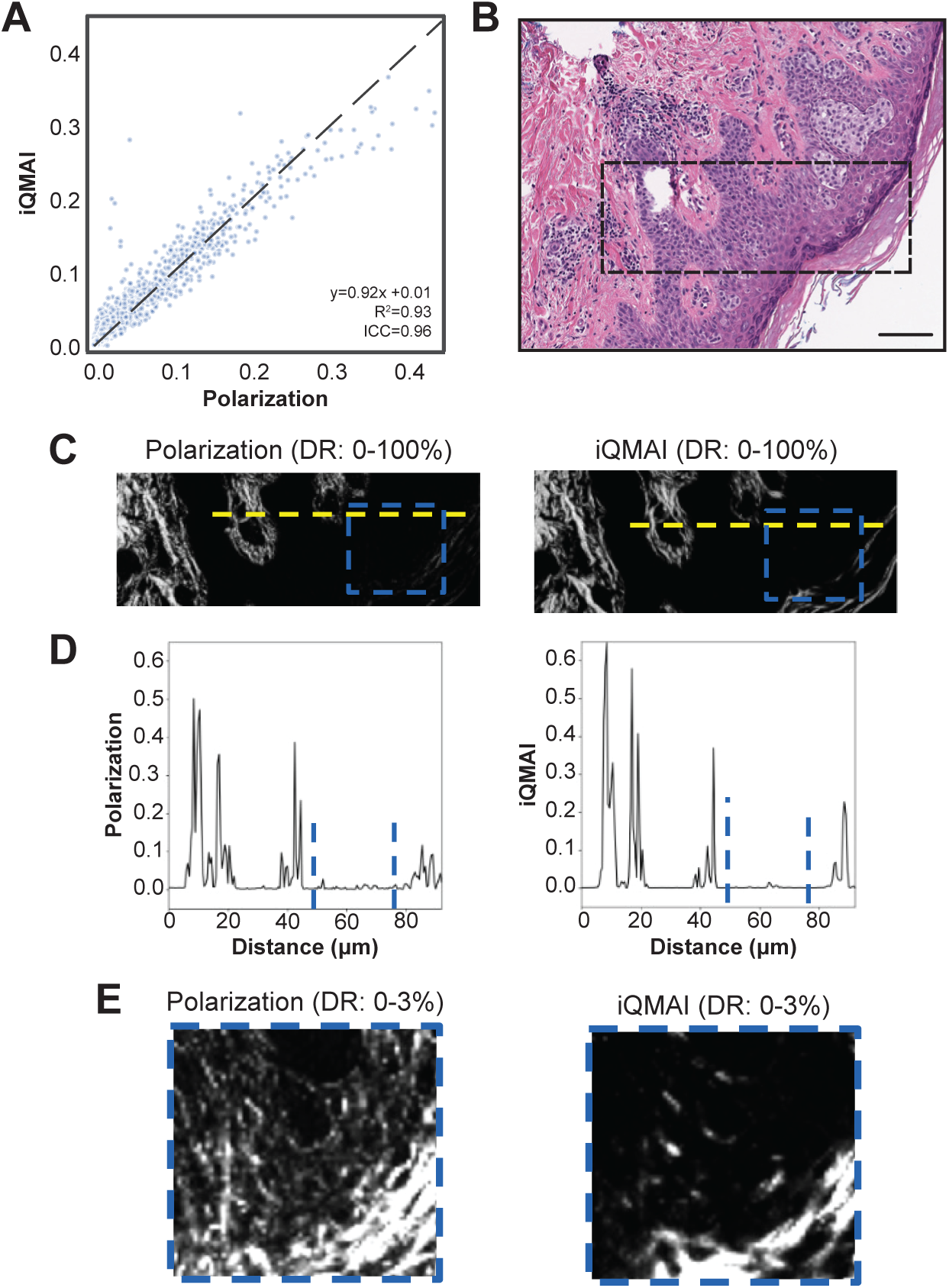
Comparison of QMAI and iQMAI outputs. A) Scatter plot of patch-wise average pixel intensities over 2293 100 μm x 100 μm image patches. B) Example region of an H&E-stained WSI used for comparison of polarization and iQMAI outputs. The region in the box was assessed by polarization and iQMAI (panel C). D) Intensity profiles over the yellow regions in panel C, showing good correspondence between polarization and iQMAI. However, the polarization intensity profile (top) was noisier in the epithelium region (between the blue dashed lines). E) Zoomed-in images of the blue regions in panels B and C clipped between 0-3% of the dynamic range. Weak scattering signals are observed in the epithelium of the QMAI image but are less prevalent in the iQMAI image. Scale bar indicates 100 μm.

The intensity profiles of images derived from ground truth polarization images and iQMAI predictions were also evaluated. As shown in an example image from melanoma (**Figure 2B**), the collagen signal (**Figure 2C**) and intensity peaks of collagen fibers were similar between the two outputs (**Figure 2D**). Notably, the signal from the ground truth image was observed to be noisier than the iQMAI signal in the tissue epithelium, a phenomenon that was due to weak scattering signals derived from epithelial cell membranes and less pronounced in iQMAI (**Figure 2E**).

In addition to collagen intensity, features describing collagen fiber morphology (fiber length, width, tortuosity, and angle relative to the epithelium-stroma interface) were extracted from ground truth and iQMAI-derived images. Image patches (300 μm x 300 μm; N=3715) were derived from the same tissue region of the ground truth and matched iQMAI images, and extracted features were directly compared, revealing a similar profile (**Figure 3A**). Direct comparison of the feature values reveals a linear relationship between those derived from ground truth images and iQMAI predictions for fiber length (R^2^=-0.17; **Figure 3B**), fiber relative angle (R^2^=0.50; **Figure 3C**), fiber width (R^2^=0.43; **Figure 3D**), and fiber tortuosity (R^2^=-0.32; **Figure 3E**). Interestingly, resulting ICC(2,1) values of median fiber width and median relative angle were significantly higher than those of median fiber length and median fiber tortuosity. Similar results were observed when assessing the agreement between features across each cancer type assessed (**Supplementary Table 4**). Further examination revealed that the performance of iQMAI was slightly diminished on short fibers that exited the focal plane, and such fibers were artificially elongated in some instances, resulting in fiber length overcall.

**Figure 3.**
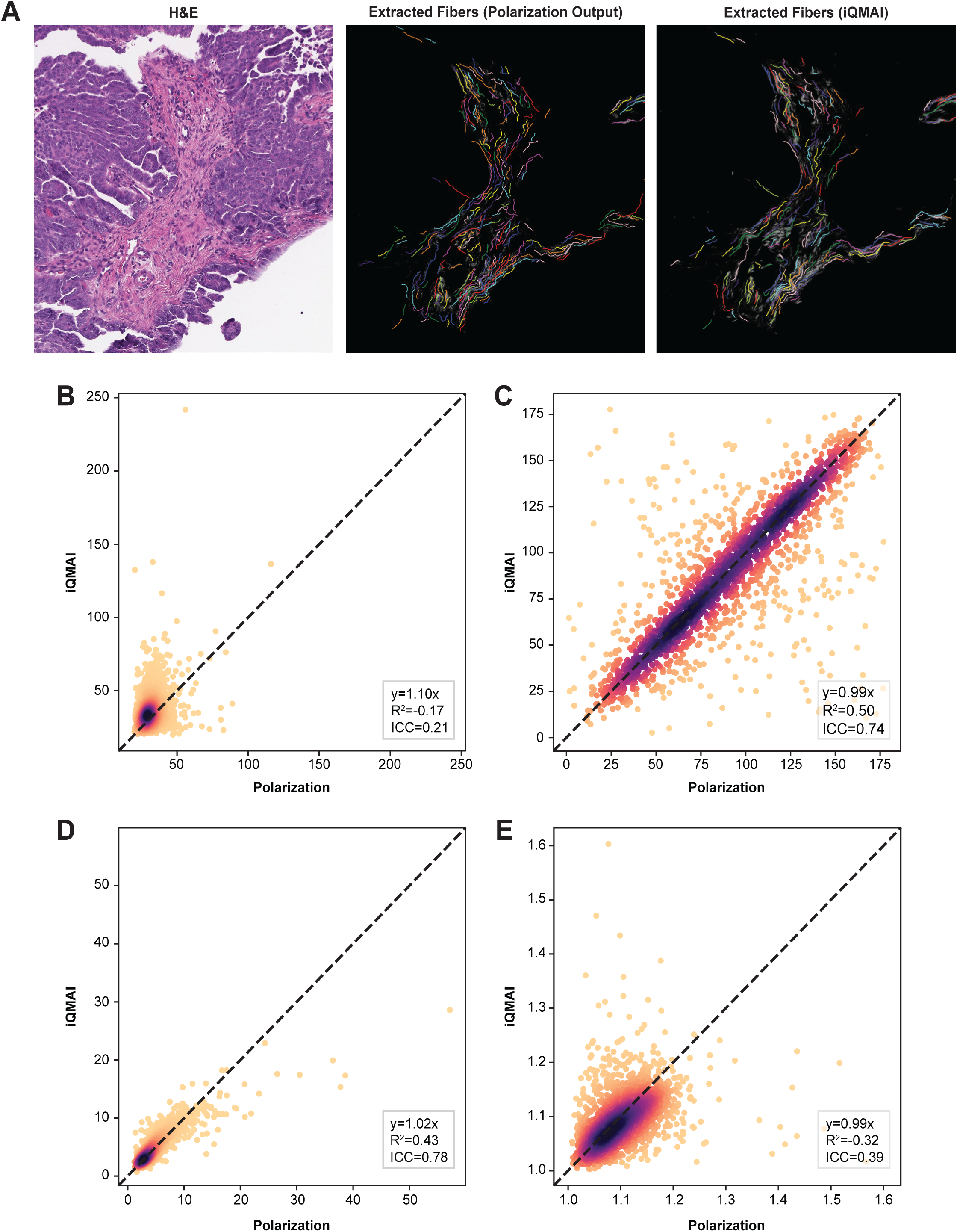
Comparison of Polarization- and iQMAI-derived fiber features. A) An example of an H&E image (left) and corresponding polarization and iQMAI collagen outputs. B-E) Scatter plots for fiber features derived from polarization and iQMAI outputs derived from 34799 patches (300 μm x 300 μm; 2000 patches selected randomly for visualization). Plots depict B) median fiber length (μm), C) median fiber relative angle, D) median fiber width (μm), and E) median fiber tortuosity Scale bars indicate 100 μm.

#### Robustness to scanner variation

Differences in imaging conditions, such as the microscope used, can lead to significant color variations between WSI. Therefore, we sought to evaluate the robustness of iQMAI to this phenomenon. H&E-stained slides (N=30) were imaged using each of four different scanning microscopes: AT2, DP200, GT450, and C13220 (**Supplementary Figure 2**). The iQMAI model was deployed on the registered H&E-stained WSIs and frames (300 μm x 300 μm; N=79670) were randomly sampled for evaluation. The iQMAI-derived collagen intensity values were compared across the four scanners and compared to values derived from AT2-scanned images. A high degree of model robustness between scanners was observed: comparisons of collagen intensity values between the AT2 and both the DP200 and C13220 yielded ICC(2,1) values of 0.92, while comparisons of such values between the AT2 and GT450 yielded an ICC(2,1) of 0.98 (**Supplementary Table 5**).

#### Pathologist evaluation

Qualitative evaluation of iQMAI was performed on randomly generated frames (N=100) from WSI (N=41) derived from 14 cancer indications. Due to the variability of collagen present in different organs, outer layers of thick muscularized blood vessels served as an internal control for collagen deposition. iQMAI showed excellent sensitivity in these vessel walls as well as in dermis of the skin and fibrous breast tissue. Neoplasms that generate a fibrotic response, such as pancreatic adenocarcinoma and cholangiocarcinoma, also showed excellent recall. Areas expected to show a dearth of collagen (i.e., lymph nodes, lymphoid tissue, and within cancer epithelium) did not show overcall of iQMAI. Similarly, indications such as DLBCL, SCLC and RCC, which are unlikely to exhibit fibrosis, did not show significant iQMAI signal.

Due to the fact that collagen can resemble smooth muscle due to its fibrous and eosinophilic texture, tissues from organs with a high degree of smooth muscle (e.g., from gastrointestinal specimens) showed minor iQMAI overcall (**Supplementary Figure 3**). Other instances of false positives occurred in regions containing other intrinsic structures known to have birefringence, such as keratin (**Supplementary Figure 4**). In addition, slight overcall was observed in red blood cells and walls of fat vacuoles (**Supplementary Figure 5**), as well as walls of thin vessels and basement membranes of colonic glands (**Supplementary Figure 6**) due to their limited presence in the training samples.

In addition, a set of frames (600 μm x 600 μm; N=100) were randomly sampled from non-excluded areas of slides in the test set. For each frame, two images – one derived from PSR-enhanced QMAI and the other derived from iQMAI – were presented to an evaluating pathologist, who evaluated the degree to which iQMAI had overcall or undercall relative to QMAI. For the one hundred frames assessed, all (100%) showed 0-10% undercall of collagen fibers, while 71 (71%) showed overcall of less than 30% of collagen fibers (**Supplementary Figure 7**).

### Biological applications of iQMAI

While algorithms to extract collagen from tissue slides have previously been utilized, these algorithms do not allow for spatial analysis of collagen in relation to other tissue components. To examine the ability of iQMAI to yield collagen features that can be spatially resolved within the tumor microenvironment (TME), we deployed iQMAI alongside PathExplore tissue and cell prediction models in H&E-stained WSI from lung adenocarcinoma (LUAD; N=492), lung squamous cell carcinoma (LUSC; N=416), hepatocellular carcinoma (HCC; N=369), and pancreatic ductal adenocarcinoma (PDAC; N=204) obtained from the Cancer Genome Atlas (TCGA).

For each cancer type examined, we compared the iQMAI-predicted density of collagen within cancer stroma to the PathExplore-predicted density of fibroblasts in the cancer stroma (**Figure 4A**). Interestingly, the distributions of collagen-fibroblast density measurements were unique for HCC, non-small cell lung cancer (NSCLC; LUAD and LUSC), and PDAC (**Figure 4B**). Collagen fiber density was correlated to a higher degree with fibroblast density in PDAC and HCC (Spearman rho=0.59 and 0.64, respectively) than in NSCLC (rho=0.32). In addition, we sought to examine collagen morphology in the stroma as a function of distance from the cancer epithelium. In regions 60, 120, and 120+ μm from the ESI, fiber width, fiber tortuosity, and relative fiber angle (relative to the ESI) were computed. In NSCLC, PDAC, and HCC, significant increases in fiber width (**Figure 4C**) and tortuosity (**Figure 4D**) were observed between regions 60 μm and 120 μm from the ESI (p<0.001 for all comparisons). Fiber angle also significantly increased between regions 60 μm and 120 μm from the ESI for NSCLC (p<0.001; **Figure 4E**), PDAC (p=0.005; **Figure 4E**), and HCC (p=0.002; **Figure 4E**). Similarly, in both NSCLC and PDAC, significant increases in fiber width (**Figure 4C**) and tortuosity (**Figure 4D**) were observed between regions 60-120 μm and >120 μm from the ESI (p<0.001 for all comparisons), while fiber angle also significantly increased between these regions (p<0.001 for NSCLC; p=0.015 for PDAC; **Figure 4E**). No significant differences in collagen fiber morphology was observed between regions 60-120 μm and >120 μm from the ESI in HCC (**Figures 4C-E**).

**Figure 4.**
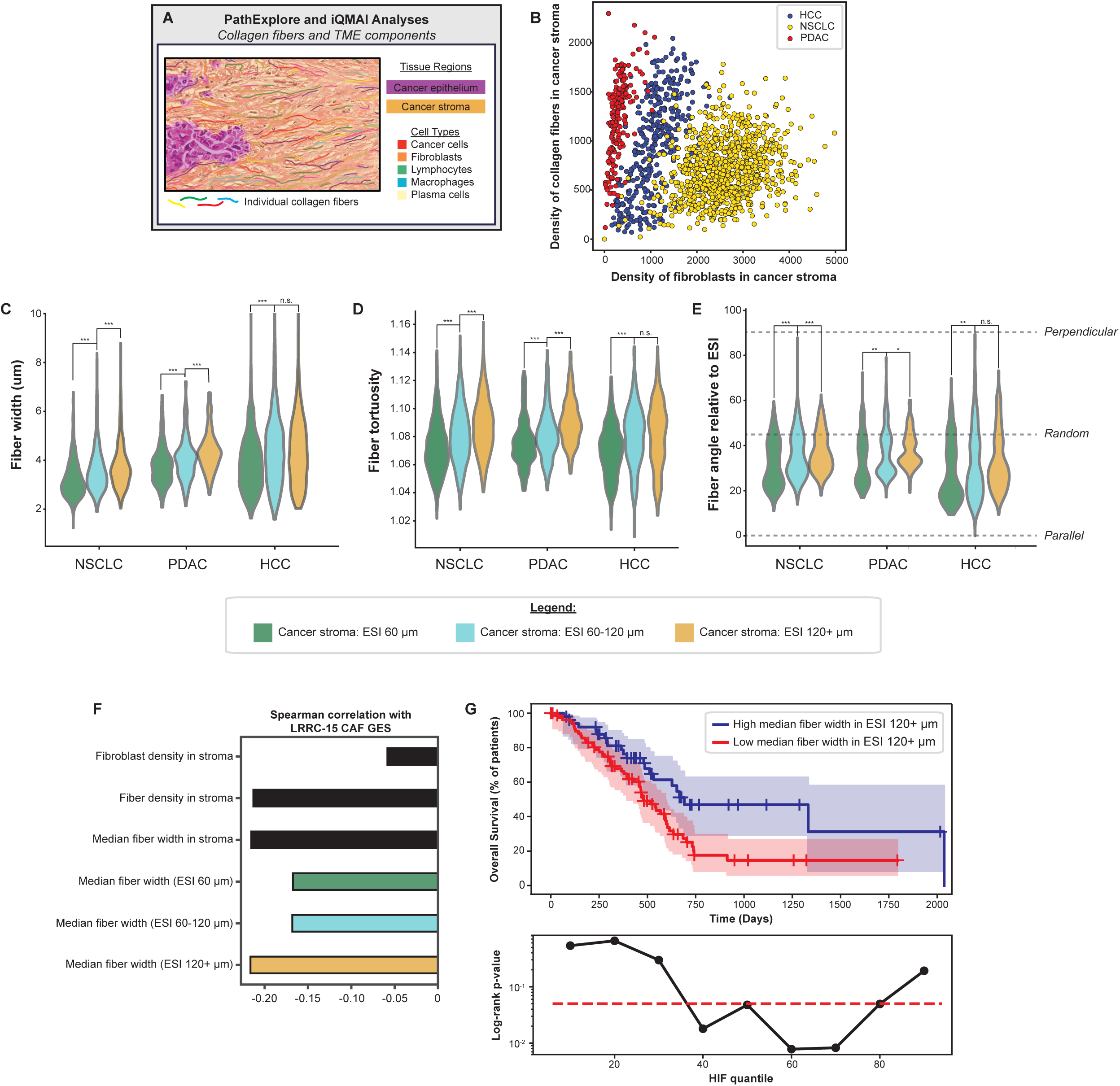
Organizational principles of collagen in cancer stroma across indications. A) iQMAI and PathExplore were simultaneously deployed to characterize collagen fibers and the TME. B) Stromal composition of different tumor types. C-E) Fiber morphology varies with distance from the epithelial-stromal interface (ESI). E) Tumors may fall into two groups based on fiber orientation relative to ESI. F) Correlation of cell and fiber features with the LRRC15 CAF signature. G) The feature showing strongest anti-correlation with the CAF signature predicts improved outcomes (HR adjusted for stage: 0.68 [0.49, 0.94]). Asterisks in panels indicate p-values calculated by Mann Whitney U test, as follows: ***: p<0.001; **: p<0.01; *: p<0.05.

In addition, we compared PathExplore-derived fibroblast features and iQMAI-derived features of collagen and individual collagen fibers to a gene expression signature known to correlate with poor response to immunotherapy, LRRC15-CAF,^47^ in PDAC (**Figure 4F**). Density of fibroblasts identified from H&E WSI did not show strong association with this signature. In contrast, overall fiber density and median fiber width were negatively correlated with this signature. Investigation of fiber features as a function of distance to the ESI revealed that the strongest association was for fibers located in the stroma beyond 120μm from the ESI. Because of the known immunosuppressive effect of LRRC15-CAFs and resulting worse prognosis, we hypothesized that fiber width in this region of the stroma might have prognostic value as well. Indeed, wider collagen fibers were associated with improved overall survival in this dataset (n=132, **Figure 4G**).

## Discussion

Altered collagen accompanies the progression of many diseases, including many cancers. However, the detection of collagen in histologic specimens has been hampered by the need for a separately stained tissue slide or a specialized imaging procedure requiring an unstained slide. Here, we used the output of a custom polarization imaging modality as ground truth to train an algorithm to infer collagen signal on digital WSIs of H&E-stained tissue.

To date, the training of ML models to detect substances in pathological specimens has traditionally required manual annotations of these substances by highly trained individuals. While this approach works well for the identification of cell types and tissue regions, which are easily distinguished by eye in a WSI, individual collagen bundles cannot be easily distinguished by visual inspection alone. As such, we sought to use collagen signals from an imaging modality for model training. In considering existing techniques for this purpose, limitations for each became apparent. Notably, SHG is limited to small regions of interest of unstained slides, which would not allow us to generate a scalable model to be used on an H&E-stained WSI. In addition, linear polarization imaging modalities are fiber orientation-dependent, while circular polarization imaging modalities are optimized for one wavelength (i.e., image color) and have suboptimal signal to noise ratio. Therefore, we elected to develop a new polarization-based imaging setup for the purpose of generating training images for this model. Images derived from this approach yielded highly accurate collagen detection on H&E- and PSR-stained tissue compared to neighboring unstained sections images with SHG. It is worthwhile to note that a model to detect collagen from an H&E image has previously been reported.^48^ That said, the prior approach used SHG signals for model training, which has several drawbacks. The SHG images used for training captured very small regions within tissue, and SHG itself requires a long imaging time and is quite expensive. Consequently, limited slides from few indications were used for model training, resulting in a less generalizable model. In contrast, the polarization imaging approach used to generate ground truth images for this work, can be applied to entire histologic specimens. Furthermore, the collagen-prediction model itself has improved architecture compared to the prior approach, incorporating a mix of perception and DICE loss and additional layers to the U-net. Therefore, while the use of microscopy images as an automated ground truth source is not novel, the improvements that we have made to this approach allow for a more accurate and generalizable collagen inference model.

The iQMAI model, itself, provides a virtual collagen overlay from H&E-stained images with high accuracy and specificity compared to polarization and SHG microscopy. Notably, an additional major advantage of the iQMAI approach is that physical glass slides and complex imaging hardware, previously required for collagen imaging, are not needed. In addition, iQMAI generalizes well on images obtained from many of the most commonly used slide scanners. Therefore, iQMAI has the potential for broad applicability across digital pathology workflows.

Another benefit of iQMAI is the potential to combine it with other analysis modalities on a H&E-stained WSI. Here, we deployed iQMAI in combination with PathExplore, a suite of ML models to comprehensively characterize tumor microenvironment,^49,50^ to understand how iQMAI-derived features can synergize with other digital pathology features. Using this approach, we not only identified an indication-specific relationship between collagen density and fibroblast density but also more general trends in how collagen morphology changes with increasing distance from the tumor epithelium. Across NSCLC, PDAC, and HCC, we observed stromal collagen fiber width, tortuosity, and relative angle to increase as the distance from the epithelial-stromal interface increases. Interestingly, across these cancer types, we observed a bimodal distribution of relative fiber angles, each of which increased with increasing distance from the cancer epithelium. Future applications of iQMAI will likely yield additional spatially-resolved insights of the interplay between stromal collagen and clinically relevant cell types, such as lymphocytes.

The collagen features derived from iQMAI also can be assessed in combination with other relevant clinical metadata, such as gene expression,clinicopathologic features and drug outcome data. Here, we observed an association between iQMAI collagen fiber density and width with the LRRC15-CAF gene expression signature^47^ in pancreatic cancer. The gene signature, itself, contains many genes involved in collagen deposition and arrangement within the ECM, as well as genes involved in the remodeling of the ECM, including *MMP11*, *COL11A1*, *CTHRC1*, and *COL12A1*.^47^ In addition, this gene signature has been associated with an immunosuppressive tumor microenvironment and poor response to immunotherapy.^18,47^ The correlations observed between iQMAI features and this gene expression signature are consistent with the previously described results and provide further mechanistic validation that the features extracted by iQMAI are accurate and biologically relevant.

Pancreatic cancer has a prominent desmospastic stroma,^51^ and myofibroblast-derived type I collagen (Col1) is a major component of the TME in this cancer type.^52–54^ Recent *in vivo* and *in vitro* work has suggested that Col1 restrains pancreatic cancer progression.^55,56^ Here, we observed that elevated collagen fiber widths – indicative of thicker collagen bundles – is associated with improved overall survival in pancreatic cancer, supporting the notion that thicker collagen can act as a barrier against tumor progression. It should be noted, however, that the relationship between collagen and prognosis seems to depend on the cancer type – for example, Col1 is associated with poor prognosis in breast cancer,^57^ and with increased metastasis in breast and prostate cancer.^58,59^ The application of iQMAI to tissues from these indications, alongside additional digital pathology models, may help to resolve the mechanism(s) behind factors related to the disparate functions of stromal collagen across cancer types.

It is worth noting that another ML model was previously described to infer SHG signal from H&E images.^48^ However, this prior approach was specific to breast and pancreas. Our goal in training iQMAI was for it to function in a pan-cancer manner. Indeed, iQMAI was trained on tissues from 15 cancer types. In addition, the utilization of perpetual loss^60^ to optimize thin fiber reconstruction and a differentiable DICE loss to guide model improvement are novel additions to the model architecture. That said, while iQMAI was trained using samples from 15 indications, the majority of slides (∼90%) were obtained from a single, high-volume pathology lab. As such, the dataset may not cover a wide spectrum of color variations in H&E slides. We addressed this potential limitation by training our U-Net with ContriMix color augmentation for improved domain generalization.^61^

In addition, there are some caveats to iQMAI signal interpretation that we noted during the course of this study. First, while iQMAI is, generally, specific in its detection of collagen, we observed that certain tissue components (e.g., fibrin and keratin) and extrinsic components (e.g., surgical ink) that harbor intrinsic birefringence had the potential to yield non-specific iQMAI signal. Minor overcall was also noted in smooth muscle and skeletal regions, both of which have eosinic fibers. As such, the best way to mitigate the overcall that arises in these nonspecific regions is to mask iQMAI signals to regions of cancer stroma, which can be identified through tissue segmentation models such as PathExplore.^50^ Another limitation of iQMAI arises due to the fact that the input images are two-dimensional slices of a three-dimensional tumor. Collagen fibers and bundles which are extracted from an H&E-stained WSI using iQMAI may enter and exit the focal plane of the image. Thus, the fiber length features that can be extracted from the iQMAI pipeline must be interpreted carefully.

In conclusion, iQMAI is a novel ML approach to detect collagen and compute collagen features from WSI of H&E-stained cancer specimens. The ability of iQMAI not only to detect collagen and quantify its morphology but also to combine iQMAI with other digital pathology models has the potential to inform the development of precision biomarkers in oncology.

## Methods

### Polarization imaging to generate model training images

An imaging modality based on the principle of polarization microscopy was developed, as follows. The imaging apparatus was built through modification of a BX63 research grade inverted microscope (Olympus Life Science, Waltham, MA) (**Supplementary Figure 8A**). Two linear polarizers (Edmund Optics #66-182 and #47-315, Barrington, NJ, USA) were inserted into the light path. The first polarizer was positioned between a switchable multi-LED illuminator (CoolLED pE-800, UK) and the specimen to generate linearly polarized incident light. The second polarizer (referred to as the analyzer) was placed in the rear objective aperture and was cross-polarized with the first polarizer. High-performance glass linear polarizers with a high extinction ratio of 10,000:1 over the range of 400-700nm were used to maximize the signal-to-noise ratio. Both polarizers were controlled by a custom-built rotational stage with a resolution of 0.028 degrees and were kept cross-polarized during rotation. Both rotational stages were controlled using a Galil DMC-4183 board.

During imaging with this approach, image stacks were acquired with multispectral polarization imaging. For multi-spectral imaging, the LED in the illuminator was switched to different wavelengths, in this case 400, 435, 470, 500, 550, 580, and 635 nm. Additional optical band pass filters were inserted in the illuminator to further narrow the full-width-half-maximum of each wavelength to approximately 10 nm. The polarizers were retracted in this mode. For polarization imaging, two intensity images were obtained at two discrete angular positions of the polarizers (45 degrees apart) and subsequently compounded to generate a polarization image. The compounding step removed the extinction zones of polarization microscopy resulting from a single angular position (**Supplementary Figure 8B**). The polarization intensity image obtained can be written as I_total_(r)=|E_i_(r)|^2^sin^2^[***Γ***(r)/2], where E_i_ is the incident electromagnetic field, r is the spatial coordinate in the sample plane, ***Γ*** is the phase shift induced by the difference in refractive indices between the fast and slow polarization axes of the specimen, and sin^2^[***Γ***(r)/2] is the quantity that contains information about the birefringence of the sample. Here, ***Γ***(r) =***β***[n_s_(r)-n_f_(r)]L is the phase shift introduced by the difference in refractive indices n_s_(r) and n_f_(r) of the slow and fast axes of the sample, respectively.

### Datasets and slide staining

Two slide sets were used. Slide set A included 337 slides from 15 indications for iQMAI development (**Supplementary Table 6**). Slides in this cohort consisted of unstained tissue microarrays (TMAs; N=38), biopsies (N=59), and surgical resections (N=240). The TMA slides were sourced from Tissue Array (Derwood, MD), while the biopsy and resection samples were from the PathAI Diagnostics (Memphis, TN) inventory in a manner to ensure a comprehensive and even representation of disease stages, histological subtypes, and procedures. Slide set B consisted of surgical resections (N=30) from 11 indications (**Supplementary Table 6**) to evaluate the model’s robustness to withstand color variation from different H&E tissue scanners.

Slides stained with hematoxylin & eosin (H&E) using using a Prisma slide stainer (Sakura, Torrance, CA) using standard conditions (**Supplementary Table 7**) and were imaged using an AT2 scanning microscope (Leica Biosystems, Vista, CA). Next, the coverslip of the H&E-stained slide was removed, and the tissue was de-stained using a solution of 0.12M HCl in ethanol, with the incubation duration dependent on the specific tissue type. The slides were then baked for fifteen minutes at 68-72°C and prior to tissue rehydration (**Supplementary Table 8**). The slides were then immersed in the PSR solution and incubated at room temperature for 60 minutes. Afterward, they were sequentially rinsed in two changes of 0.5% acetic acid for 25 seconds each, followed by rinsing in 100% ethanol. Finally, the slides were left to air-dry in preparation for imaging.

The PSR-stained slide was then imaged using the polarization workflow, acquiring one polarization image at 635 nm and one MSI image at 435 nm (**Figure 5C**). An additional image registration step was used to register the H&E image against the polarization image by estimating the registration mapping between the MSI image and the H&E image. All polarization images used in the study had an exposure time of 3.75 ms. This process was repeated for each FOV to ensure full coverage of the WSI (**Figure 5D**).

**Figure 5.**
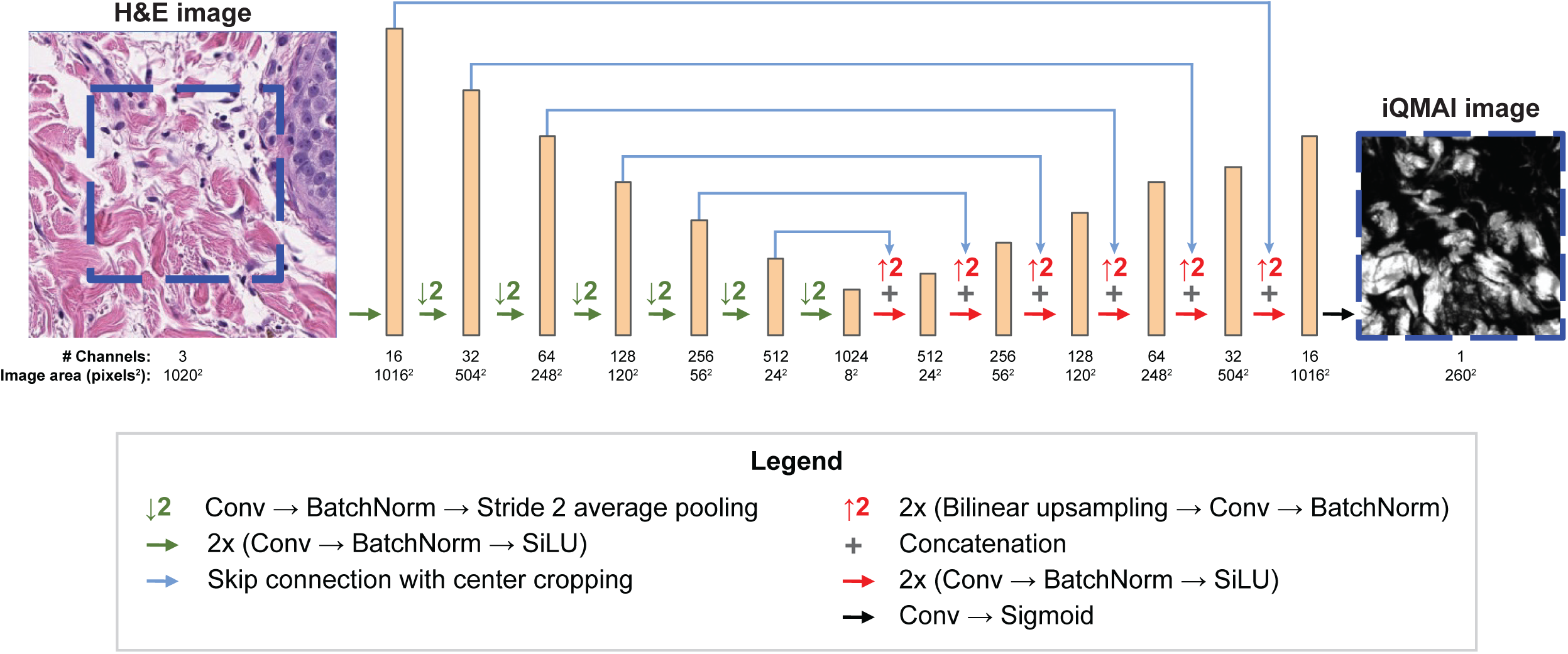
iQMAI architecture. The U-Net architecture used for H&E to iQMAI image translation.

### Second Harmonic Generation

Neighboring unstained sections were imaged using a Leica TCS SP8 microscope. The system used a femtosecond-pulsed laser (Insight DS+) with an adjustable wavelength within 680-1300nm. An excitation wavelength of 860nm was used with a 2 percent laser power intensity (1.4W). The field-of-view consisted of 512 x 512 pixels with a pixel size of 0.91 μm. Throughout the scanning process, the Fluotar VISIR 25x/0.95 water immersion objective was utilized. A 488nm filter (20 nm bandwidth) was placed before the photomultiplier tube of the scanner to block non-second harmonic signals. Additionally, line averaging was performed over eight consecutive scans to enhance the signal-to-noise ratio of SHG. The raw output images, in .lif 16-bit format, were converted to 8-bit PNG format for comparison with polarization images.

### Quality control

After data acquisition, a strict data quality control (QC) pipeline identified unsatisfactory regions to be excluded from the study. A board-certified pathologist carefully inspected all images of the PSR-stained slides to identify suggested regions to be excluded due to poor staining, poor focus, scanning errors, missing tissue from re-staining, and regions missed during scanning. In addition, an automatic missing tissue QC process utilized image processing to identify missing tissue by comparing the H&E image to the MSI image. In short, the input H&E WSI was converted to grayscale, which was transformed into a binary image using Otsu’s thresholding. Subsequently, background pixels identified from the binarized WSI image were used to calculate the average background pixel intensity in the pre-registered MSI image, which served as a threshold to binarize the MSI image. The disparity between the segmented grayscale WSI and the segmented MSI produced a missing tissue binary mask. This mask was combined with another binary mask that identifies regions not scanned by the polarization system, resulting in a mask of areas to be excluded.

### iQMAI model architecture

A U-Net architecture^62^ was used for dense-regression to predict the iQMAI signal from the input H&E image (**Figure 5**). The network consisted of a downsampling and an upsampling path with 6 levels each. Each downsampling level consisted of a 2x downsampling block with average pooling followed by a ConvNormAct block repeated twice. The ConvNormAct block serialized 2D convolution, batch normalization, and SiLU activation. Each upsampling level used 2x bilinear upsampling, followed by concatenating of a skipped signal from the downsampling path. The concatenated signal was passed to a ConvNormAct block to compute the output signal for the next level. A convolution block followed by Sigmoid activation was added to the output of the upsampling path to generate the output iQMAI image with the signal range in [0,1). The input patch size was 1020 x 1020 pixels with three channels, representing the H&E RGB image; the size of the output iQMAI image was 260 x 260 pixels, representing the predicted intensity of the polarization signal. Only valid convolution with no padding was used in the U-Net to avoid potential artifacts at patch boundaries.

### iQMAI model training

Slide cohorts were divided into training, validation, and test splits in a manner that maximized each split’s diversity. Slides from set A were stratified across different indications to have approximately 70% slides for training (N=244), 15% slides for validation (N=47), and 15% slides for testing (N=47). To overcome a limited number of slides from small cell lung cancer, diffuse large B cell lymphoma, and ovarian cancer specimens, 37 TMA slides were partitioned into non-overlapping regions for training (75%) and validation/testing (25%). We used the training regions within these slides for the training splits while reserving the held-out one for the validation and test splits. The number of slides for each indication per split and sampling results are shown in **Supplementary Table 9** and **Supplementary Table 10**, respectively.

For model training, pairs of registered H&E-stained and PSR-enhanced polarization frames (1020 x 1020 pixels) were extracted and randomly sampled on a rectangular grid. A significant imbalance in the usable tissue areas across different indications was observed, such as a 13-fold difference in tissue area between the indication with the largest usable area (prostate cancer) and the indication with the smallest usable area (small cell lung carcinoma). To account for this disparity, dynamic adjustments were made to the grid spacing based on the tissue area, with a larger grid spacing for indications with greater usable tissue area and vice versa, this equivalent representation of tissue areas across all indications. The minimum value of the grid spacing was limited to 25 μm to minimize similarity between training samples. To improve the sample quality, the relative shift between the MSI frame, which co-aligned to the PSR-enhanced polarization frame, to the H&E frame was estimated using template matching based on normalized cross-correlation,^63^ and all frames whose shifts were more than 5 pixels or correlation coefficient less than 0.7 were rejected. This approach yielded 4,926,844 training pairs and 666,095 validation pairs, with approximately 301,000-366,000 training pairs per indication.

A loss function was minimized over minibatch training samples between the ground truth PSR-enhanced polarization image (I_gt_) and the predicted iQMAI image (Î=f(I_HE_)) from the input H&E image (I_HE_), as follows: *ɭ*(Î, I_gt_)=**γ***ɭ_cons_*(Î, I_gt_)+(1-**γ**)*ɭ_DICE_*(Î, I_gt_;***τ***). ***τ*** was the binarization parameter that was sampled from the interval [0.0, 0.4] during training. *ɭ*(.) was the loss function, consisting of an image consistency loss component (*ɭ_cons_)* and customized DICE loss (*ɭ_DICE_*) specifically designed to address undercalls in regions with strong QMAI signal. Different reconstruction losses were used, including the mean absolute error (MAE) loss, L_MAE_(Î, I_gt_)=||Î - I_gt_||^1^/N, mean squared error (MSE) loss L_MSE_(Î, I_gt_)=||Î - I_gt_||^2^_2_/N, LPIPS only loss *ɭ_LPIPS_*(Î, I_gt_), and a combination of LPIPS and DICE loss 0.01*ɭ_LPIPS_*(Î, I_gt_) + 0.99*ɭ_DICE_*(Î, I_gt_). Using only MAE and MSE loss resulted in lower resolution, while using only LPIPS resulted in collagen fiber undercall (**Supplementary Figure 1**). Therefore, a model combining LPIPS loss with DICE loss was selected.

Finally, **γ** was a hyper-parameter that controlled the trade-off between two loss components. An investigation of the different forms of image consistency loss revealed that the perpetual loss^60^ maximized reconstruction quality and was resilient to slight misregistration between H&E and polarization training images (**Supplementary Figure 2**).

U-Nets were trained on 32 RTX 3080 GPUs on Pytorch using the AdamW optimizer^64^ with a learning rate of 5e-4 for 500,000 iterations and a minibatch size of 2560. The learning rate was decayed by a factor of 0.9 after every 10,000 iterations. An image transformation pipeline augmented each training with random horizontal and vertical flipping, random image rotation by multiples of 90 degrees, and random cropping, before passing to ContriMix stain color augmentation^61^ to generate three variations of the input H&E images. Additionally, we applied random light perturbation, additive corruption, and frequency corruption to these variations to enhance model robustness to common artifacts in H&E images. Based on an ablation experiment (**Supplementary Table 1**), a trade-off factor **γ**=0.01 was chosen.

### Custom differentiable DICE loss

A customized DICE loss was incorporated to encourage consistency between the continuous QMAI and iQMAI. The DICE loss was defined as *ɭ_DICE_*(Î, I_gt_;***τ***), where Î and I_gt_ represent the iQMAI image and the PSR-enhanced QMAI ground truth image, respectively. Both images had continuous pixel values in the range of [0.0, 1.0]. ***τ*** was the threshold parameter for binarization, randomly sampled in the interval of [0.0, 0.4] during training.

The DICE loss can be written as *ɭ_DICE_*(Î, I_gt_;***τ***)= 1 - (2||B_pred;***τ***_ B_gt;***τ***_||_1_)/(||B_pred;***τ***_||_1_+||B_gt;***τ***_||_1_), where *B_gt;_*_***τ***_=[*I_gt_ ≥* ***τ***] is the binarized iQMAI image. To make the DICE loss fully differentiable, *B_pred;_*_***τ***_ was defined as a proxy for the true binarized iQMAI image ([Î *≥* ***τ***]) as *B_pred;_*_***τ***_ = *Sigmoid*[(Î - ***τ***)/**α**], where **α** was a small temperature control parameter, defining the speed of the saturation when the pixel values in Î deviated from the threshold ***τ***. ||x||_1_ denoted the l1-norm of the image x (i.e., the summation of absolute pixel values).

### Fiber and fiber-based feature extraction

From the iQMAI signal, individual fibers and their features (e.g. width, length) were extracted using a modified FIbeR Extraction (FIRE) algorithm^65^ (modified to incorporate skeletonization and graph post-processing to merge nodes and edges to build graphs). Given the importance of pre-processing an image prior to FIRE application,^44^ a Gaussian filter was applied to the image before binarization and application of the distance transform. In order to find the nucleation points, a ridge filter^66^ was applied before the image was binarized and subsequently skeletonized. For each nucleation point location, the pixel value in the distance transformed image corresponded to the distance to the edge of the fiber. After carrying out those pre-processing steps, a python implementation of the FIRE algorithm was applied to extract the individual fibers and their features. After extracting the fibers and their individual features, aggregate fiber features at the patch- and WSI-level were derived (e.g., count of fibers or standard deviation, mean or median of fiber width or length).

### Evaluation of model robustness to scanner

H&E stained slides (N=30) from set B were imaged using four distinct scanners: AT2 (Leica Biosystems, Vista, CA), DP200 (Roche Diagnostics, Tucson, AZ), GT450 (Leica Biosystems, Vista, CA), and C13220 (Hamamatsu Photonics, Bridgewater, NJ), resulting in 4 WSIs per slide. The WSIs were then aligned to the WSI from AT2 as the reference. Five slides were excluded due to poor image registration. Following image alignment, iQMAI was deployed on the registered WSIs, and rectangular patches were extracted for intra-class correlation analysis. WSI registration occurred in two sequential steps: 1) coarse registration, which consisted of finding the affine spatial transformations (i.e. the scale, rotation, translation parameters) that coarsely align the moving WSIs to the reference WSIs at low resolution, and 2) patch-wise rigid registration at higher resolution which consisted of iterating over the patches of the coarsely aligned WSIs and finding the affine spatial transformations (i.e. rotation, translation parameters) that further aligned the patches from the moving WSIs to the patches of the reference WSIs.

Determination of the scale, rotation, and translation parameters of the coarse affine registration proceeded as follows. Microns per pixel (MPP) from the WSI metadata provided scale information and was utilized to determine the scaling factor between the AT2 WSI and corresponding WSI from the DP200, GT450, and C13220 scanners. For rotation and translation, the radon transform was implemented on a low resolution tissue binary mask (8 MPP) obtained by the deployment of an internal tissue detection model. Implementation of cross correlation on the radon-transformed image was utilized to identify the optimal rotation angle at this resolution.^67^ The rotation parameter was refined by iteratively rotating the moving image at higher resolution (1 MPP) by angles between ***θ***_low res_-1 and ***θ***_low res_+1 (in degrees) with a step of 0.1 degree, followed by a cross-correlation between the rotated moving image and the reference image to estimate the optimal translation for each angle.^68^ The rotation angle and translation corresponding to the lowest translation invariant root mean squared error were selected for the coarse affine registration. The spatial transformation derived through this process was then applied to the iQMAI images derived from each scanner, to register them to each other and enable local comparisons of the iQMAI signal across scanners.

For a more precise alignment, the moving WSI was warped using the coarse rigid transformation found in the coarse alignment step. And, iterating over the patches of both the reference WSI and coarsely aligned moving WSI at high resolution (0.5 MPP) for each pair of reference and moving patches, a rigid transformation (i.e. the rotation and translation parameters) for registration was identified by iteratively rotating the patch from the warped moving image by angles between −1 and 1 degrees with a step of 0.1 degree, followed by cross-correlation to estimate the optimal translation for each angle. The rotation angle and translation corresponding to the lowest translation invariant root mean squared error were selected as rigid registration parameters for each pair of patches.

The final spatial transformation that aligned the moving WSI to the reference WSI was the composition of the coarse WSI affine transformation and the patch-wise rigid transformations. These spatial transformations were then applied to the iQMAI images, enabling local comparison of the iQMAI signal across the AT2, DP200, GT450, and C13220 scanners.

### Analysis of collagen features in TCGA samples

H&E-stained WSI were obtained from The Cancer Genome Atlas^69^ as follows: LUAD and LUSC (NSCLC; N=908), LIHC (HCC; N=369), and PAAD (PDAC; N=204). PathExplore^TM^ (Boston, MA), a suite of ML models to segment tissue regions (cancer epithelium, cancer-associated stroma, necrosis and normal tissue),^50^ detect and classify cell types (e.g., cancer cells, lymphocytes, and fibroblasts),^49^ and segment the cancer stroma into regions <60, 60-120, and 120+ μm from the epithelial-stromal interface (ESI) were also measured. PathExplore is for research use only and is not for use in diagnostic procedures.Human interpretable features (HIFs) were subsequently extracted from WSI as previously described.^70^ In short, HIFs were generated through combining model heatmap overlays to yield the density of fibroblasts in cancer-associated stroma.

iQMAI was then deployed on the same WSI, and HIFs quantifying collagen fiber abundance and morphology were extracted as previously described.^44^ iQMAI-derived collagen overlays were combined with PathExplore-derived tissue and cell overlays to yield spatially-resolved collagen features, such as the density of collagen fibers in cancer-associated stroma and fiber width in stroma regions 60-120 μm from the ESI.

### Statistical analysis

Collagen feature values derived from polarization ground truth imaging and iQMAI were compared using linear regression using the Huber regressor,^71^ and intraclass correlation coefficient (ICC) (2,1) values were estimated. The prognostic value of fiber width was assessed by fitting a Cox proportional hazard model to overall survival duration in the TCGA PAAD cohort (N=132) using tumor stage as a covariate. For visualization using a KM curve the population was split into high/low feature values at the 60th percentile. A sensitivity analysis to this cutoff was performed by computing the log-rank p-value from the 10th to the 90th percentiles in 10 percent increments.

## References

1. Cox, T. R. The matrix in cancer. Nat. Rev. Cancer 21, 217–238 (2021).

2. Ricard-Blum, S. The collagen family. Cold Spring Harb. Perspect. Biol. 3, a004978 (2011).

3. Provenzano, P. P. et al. Collagen reorganization at the tumor-stromal interface facilitates local invasion. BMC Med. 4, 38 (2006).

4. Amatangelo, M. D., Bassi, D. E., Klein-Szanto, A. J. P. & Cukierman, E. Stroma-derived three-dimensional matrices are necessary and sufficient to promote desmoplastic differentiation of normal fibroblasts. Am. J. Pathol. 167, 475–488 (2005).

5. Mierke, C. T. Mechanical cues affect migration and invasion of cells from three different directions. Front. Cell Dev. Biol. 8, 583226 (2020).

6. Conklin, M. W. et al. Aligned collagen is a prognostic signature for survival in human breast carcinoma. Am. J. Pathol. 178, 1221–1232 (2011).

7. Conklin, M. W. et al. Collagen alignment as a predictor of recurrence after ductal carcinoma in situ. Cancer Epidemiol. Biomarkers Prev. 27, 138–145 (2018).

8. Esbona, K. et al. The presence of cyclooxygenase 2, tumor-associated macrophages, and collagen alignment as prognostic markers for invasive breast carcinoma patients. Am. J. Pathol. 188, 559–573 (2018).

9. Hanley, C. J. et al. A subset of myofibroblastic cancer-associated fibroblasts regulate collagen fiber elongation, which is prognostic in multiple cancers. Oncotarget 7, 6159–6174 (2016).

10. Fanous, M., Keikhosravi, A., Kajdacsy-Balla, A., Eliceiri, K. W. & Popescu, G. Quantitative phase imaging of stromal prognostic markers in pancreatic ductal adenocarcinoma. Biomed. Opt. Express 11, 1354–1364 (2020).

11. Almici, E. et al. Quantitative image analysis of fibrillar collagens reveals novel diagnostic and prognostic biomarkers and histotype-dependent aberrant mechanobiology in lung cancer. Mod. Pathol. 36, 100155 (2023).

12. Sadjadi, Z., Zhao, R., Hoth, M., Qu, B. & Rieger, H. Migration of cytotoxic T lymphocytes in 3D collagen matrices. Biophys. J. 119, 2141–2152 (2020).

13. Wolf, K. et al. Physical limits of cell migration: control by ECM space and nuclear deformation and tuning by proteolysis and traction force. J. Cell Biol. 201, 1069–1084 (2013).

14. Pruitt, H. C. et al. Collagen fiber structure guides 3D motility of cytotoxic T lymphocytes. Matrix Biol. 85-86, 147–159 (2020).

15. Kuczek, D. E. et al. Collagen density regulates the activity of tumor-infiltrating T cells. J. Immunother. Cancer 7, (2019).

16. Xiao, Z. et al. Desmoplastic stroma restricts T cell extravasation and mediates immune exclusion and immunosuppression in solid tumors. Nat. Commun. 14, 5110 (2023).

17. Peng, D. H. et al. Collagen promotes anti-PD-1/PD-L1 resistance in cancer through LAIR1-dependent CD8+ T cell exhaustion. Nat. Commun. 11, 4520 (2020).

18. Chakravarthy, A., Khan, L., Bensler, N. P., Bose, P. & De Carvalho, D. D. TGF-β-associated extracellular matrix genes link cancer-associated fibroblasts to immune evasion and immunotherapy failure. Nat. Commun. 9, 4692 (2018).

19. Garvey, W., Fathi, A., Bigelow, F., Carpenter, B. & Jimenez, C. Improved Movat pentachrome stain. Stain Technol. 61, 60–62 (1986).

20. Foot, N. C. The Masson Trichrome Staining Methods in Routine Laboratory Use. Stain Technol. 8, 101–110 (1933).

21. Whittaker, P. & Canham, P. B. Demonstration of quantitative fabric analysis of tendon collagen using two-dimensional polarized light microscopy. Matrix 11, 56–62 (1991).

22. Yakovlev, D. D. et al. Quantitative mapping of collagen fiber alignment in thick tissue samples using transmission polarized-light microscopy. J. Biomed. Opt. 21, 71111 (2016).

23. Spiesz, E. M., Kaminsky, W. & Zysset, P. K. A quantitative collagen fibers orientation assessment using birefringence measurements: calibration and application to human osteons. J. Struct. Biol. 176, 302–306 (2011).

24. Lattouf, R. et al. Picrosirius red staining: a useful tool to appraise collagen networks in normal and pathological tissues: A useful tool to appraise collagen networks in normal and pathological tissues. J. Histochem. Cytochem. 62, 751–758 (2014).

25. Junqueira, L. C., Bignolas, G. & Brentani, R. R. Picrosirius staining plus polarization microscopy, a specific method for collagen detection in tissue sections. Histochem. J. 11, 447–455 (1979).

26. Drifka, C. R. et al. Comparison of picrosirius red staining with second harmonic generation imaging for the quantification of clinically relevant collagen fiber features in histopathology samples. J. Histochem. Cytochem. 64, 519–529 (2016).

27. Williams, R. M., Zipfel, W. R. & Webb, W. W. Interpreting second-harmonic generation images of collagen I fibrils. Biophys. J. 88, 1377–1386 (2005).

28. Chen, X., Nadiarynkh, O., Plotnikov, S. & Campagnola, P. J. Second harmonic generation microscopy for quantitative analysis of collagen fibrillar structure. Nat. Protoc. 7, 654–669 (2012).

29. Keikhosravi, A., Bredfeldt, J. S., Sagar, A. K. & Eliceiri, K. W. Second-harmonic generation imaging of cancer. Methods Cell Biol. 123, 531–546 (2014).

30. Módis, L. Organization of the Extracellular Matrix: A Polarization Microscopic Approach. (CRC Press, 2018).

31. Keikhosravi, A. et al. Real-time polarization microscopy of fibrillar collagen in histopathology. Sci. Rep. 11, 19063 (2021).

32. Chue-Sang, J. et al. Use of Mueller matrix colposcopy in the characterization of cervical collagen anisotropy. J. Biomed. Opt. 23, 1–9 (2018).

33. Tumanova, K. et al. Mueller matrix polarization parameters correlate with local recurrence in patients with stage III colorectal cancer. Sci. Rep. 13, 13424 (2023).

34. Mirsanaye, K. et al. Machine learning-enabled cancer diagnostics with widefield polarimetric second-harmonic generation microscopy. Sci. Rep. 12, 10290 (2022).

35. Dong, J., Yao, Y., Dong, Y. & Ma, H. From pixel to slide image: Polarization modality-based pathological diagnosis using representation learning. arXiv [eess.IV] (2024).

36. Huang, T. et al. Mueller matrix imaging of pathological slides with plastic coverslips. Opt. Express 31, 15682–15696 (2023).

37. Dong, Y. et al. A polarization-imaging-based machine learning framework for quantitative pathological diagnosis of cervical precancerous lesions. IEEE Trans. Med. Imaging 40, 3728–3738 (2021).

38. Majumdar, A. et al. Machine learning based local recurrence prediction in colorectal cancer using polarized light imaging. J. Biomed. Opt. 29, 052915 (2024).

39. Lad, J., Serra, S., Quereshy, F., Khorasani, M. & Vitkin, A. Polarimetric biomarkers of peri-tumoral stroma can correlate with 5-year survival in patients with left-sided colorectal cancer. Sci. Rep. 12, 12652 (2022).

40. Deng, L. et al. Influence of hematoxylin and eosin staining on linear birefringence measurement of fibrous tissue structures in polarization microscopy. J. Biomed. Opt. 28, 102909 (2023).

41. Yao, Y. et al. Polarization imaging-based radiomics approach for the staging of liver fibrosis. Biomed. Opt. Express 13, 1564–1580 (2022).

42. Mi, C., Shao, C., He, H., He, C. & Ma, H. Evaluating tissue mechanical properties using quantitative Mueller matrix polarimetry and neural network. Appl. Sci. (Basel) 12, 9774 (2022).

43. Shi, Y., Chen, B., He, H. & Ma, H. Quantitative assessment of tissue structures based on Mueller matrix polarimetry and derived parameters imaging. in Polarized Light and Optical Angular Momentum for Biomedical Diagnostics 2022 (eds. Ramella-Roman, J. C., Ma, H., Vitkin, I. A., Elson, D. S. & Novikova, T.) (SPIE, 2022). doi:10.1117/12.2609364.

44. Bredfeldt, J. S. et al. Computational segmentation of collagen fibers from second-harmonic generation images of breast cancer. J. Biomed. Opt. 19, 16007 (2014).

45. Rittié, L. Method for picrosirius red-polarization detection of collagen fibers in tissue sections. Methods Mol. Biol. 1627, 395–407 (2017).

46. Bakurov, I., Buzzelli, M., Schettini, R., Castelli, M. & Vanneschi, L. Structural similarity index (SSIM) revisited: A data-driven approach. Expert Syst. Appl. 189, 116087 (2022).

47. Dominguez, C. X. et al. Single-Cell RNA Sequencing Reveals Stromal Evolution into LRRC15+ Myofibroblasts as a Determinant of Patient Response to Cancer Immunotherapy. Cancer Discov. 10, 232–253 (2020).

48. Keikhosravi, A. et al. Non-disruptive collagen characterization in clinical histopathology using cross-modality image synthesis. Commun. Biol. 3, 414 (2020).

49. Abel, J. et al. AI powered quantification of nuclear morphology in cancers enables prediction of genome instability and prognosis. npj Precision Oncology 8, 134 (2024).

50. Markey, M., et al. Spatial mapping of immunosuppressive CAF gene signatures in H&E-stained images using additive multiple instance learning. bioRxiv (2024) doi:10.1101/2024.08.12.607604.

51. Mahadevan, D. & Von Hoff, D. D. Tumor-stroma interactions in pancreatic ductal adenocarcinoma. Mol. Cancer Ther. 6, 1186–1197 (2007).

52. Mollenhauer, J., Roether, I. & Kern, H. F. Distribution of extracellular matrix proteins in pancreatic ductal adenocarcinoma and its influence on tumor cell proliferation in vitro. Pancreas 2, 14–24 (1987).

53. Imamura, T. et al. Quantitative analysis of collagen and collagen subtypes I, III, and V in human pancreatic cancer, tumor-associated chronic pancreatitis, and alcoholic chronic pancreatitis. Pancreas 11, 357–364 (1995).

54. Tian, C. et al. Proteomic analyses of ECM during pancreatic ductal adenocarcinoma progression reveal different contributions by tumor and stromal cells. Proc. Natl. Acad. Sci. U. S. A. 116, 19609– 19618 (2019).

55. Chen, Y. et al. Type I collagen deletion in αSMA+ myofibroblasts augments immune suppression and accelerates progression of pancreatic cancer. Cancer Cell 39, 548–565.e6 (2021).

56. Chen, Y. et al. Oncogenic collagen I homotrimers from cancer cells bind to α3β1 integrin and impact tumor microbiome and immunity to promote pancreatic cancer. Cancer Cell 40, 818–834.e9 (2022).

57. Barcus, C. E. et al. Elevated collagen-I augments tumor progressive signals, intravasation and metastasis of prolactin-induced estrogen receptor alpha positive mammary tumor cells. Breast Cancer Res. 19, 9 (2017).

58. Kakkad, S. M. et al. Collagen I fiber density increases in lymph node positive breast cancers: pilot study. J. Biomed. Opt. 17, 116017 (2012).

59. Penet, M.-F. et al. Structure and function of a prostate cancer dissemination-permissive extracellular matrix. Clin. Cancer Res. 23, 2245–2254 (2017).

60. Zhang, R., Isola, P., Efros, A. A., Shechtman, E. & Wang, O. The unreasonable effectiveness of deep features as a perceptual metric. arXiv [cs.CV] (2018).

61. Nguyen, T. H., et al. ContriMix: Scalable stain color augmentation for domain generalization without domain labels in digital pathology. arXiv [eess.IV] (2023).

62. Ronneberger, O., Fischer, P. & Brox, T. U-Net: Convolutional Networks for Biomedical Image Segmentation. arXiv [cs.CV] (2015).

63. Briechle, K. & Hanebeck, U. D. Template matching using fast normalized cross correlation. in Optical Pattern Recognition XII (eds. Casasent, D. P. & Chao, T.-H.) (SPIE, 2001). doi:10.1117/12.421129.

64. Loshchilov, I. & Hutter, F. Decoupled weight decay regularization. arXiv [cs.LG] (2017).

65. Stein, A. M., Vader, D. A., Jawerth, L. M., Weitz, D. A. & Sander, L. M. An algorithm for extracting the network geometry of three-dimensional collagen gels. J. Microsc. 232, 463–475 (2008).

66. Frangi, A. F., Niessen, W. J., Vincken, K. L. & Viergever, M. A. Multiscale vessel enhancement filtering. in Medical Image Computing and Computer-Assisted Intervention — MICCAI’98 130–137 (Springer Berlin Heidelberg, Berlin, Heidelberg, 1998).

67. Chelbi, S. & Mekhmoukh, A. Features based image registration using cross correlation and Radon transform. Alex. Eng. J. 57, 2313–2318 (2018).

68. Guizar-Sicairos, M., Thurman, S. T. & Fienup, J. R. Efficient subpixel image registration algorithms. Opt. Lett. 33, 156–158 (2008).

69. Gutman, D. A. et al. Cancer Digital Slide Archive: an informatics resource to support integrated in silico analysis of TCGA pathology data. J. Am. Med. Inform. Assoc. 20, 1091–1098 (2013).

70. Diao, J. A. et al. Human-interpretable image features derived from densely mapped cancer pathology slides predict diverse molecular phenotypes. Nat. Commun. 12, 1613 (2021).

71. Huber, P. J. Robust estimation of a location parameter. in Springer Series in Statistics 492–518 (Springer New York, New York, NY, 1992).

