## Supplemental Figures and Tables for "Quantification of collagen and associated features from H&E-stained whole slide pathology images across cancer types using a physics-based deep learning model"

### Supplementary Figures

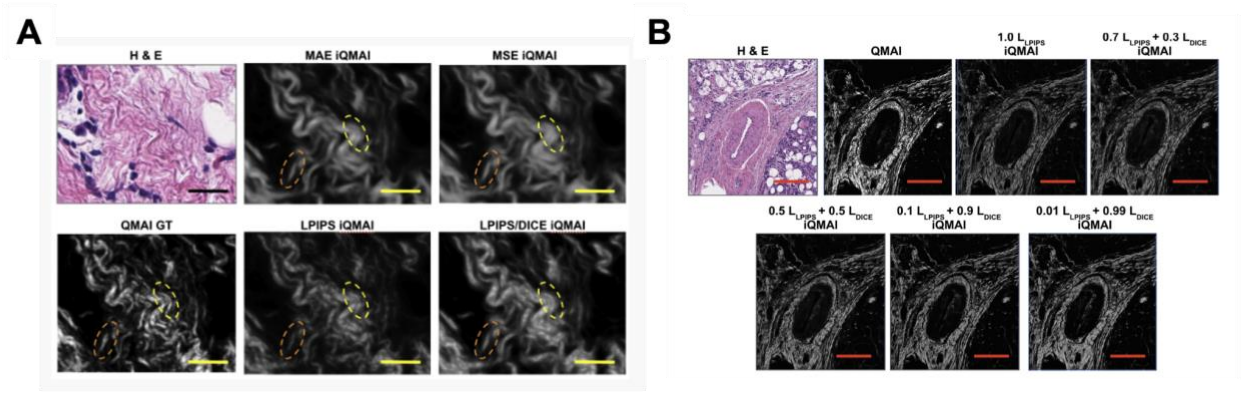


**Supplementary Figure 1.** A) Visualization of iQMAI pan-cancer results trained with different reconstruction loss functions. Scale bars indicate 25 μm. B) Visualization of iQMAI pan-cancer results from models trained with different combination of $\mathcal{L}_{LPIPS}$ and $\mathcal{L}_{DICE}$. Scale bars indicate 200 μm.

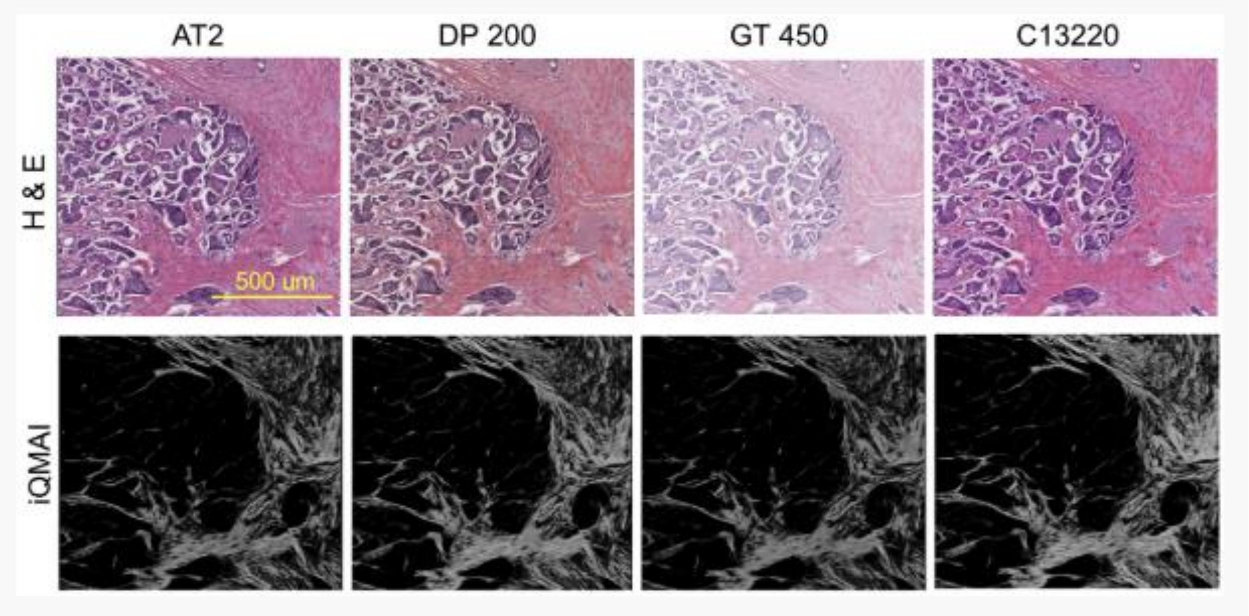


**Supplementary Figure 2. Robustness of iQMAI to scanner variability.** Example H&E images from 4 different scanners (top) and their corresponding iQMAI images (bottom) on a breast sample.

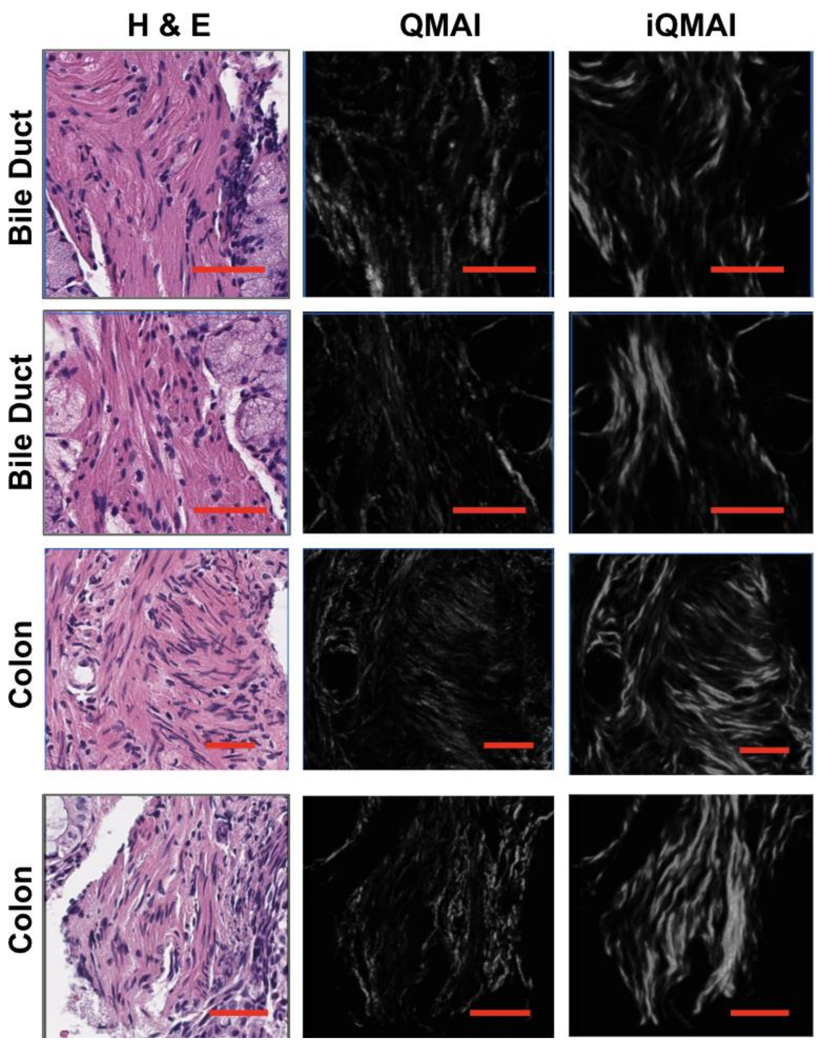


**Supplementary Figure 3.** iQMAI overcall in smooth muscle of the bile duct (top 2 rows) and colons (bottom 2 rows). Scale bar, 50 μm

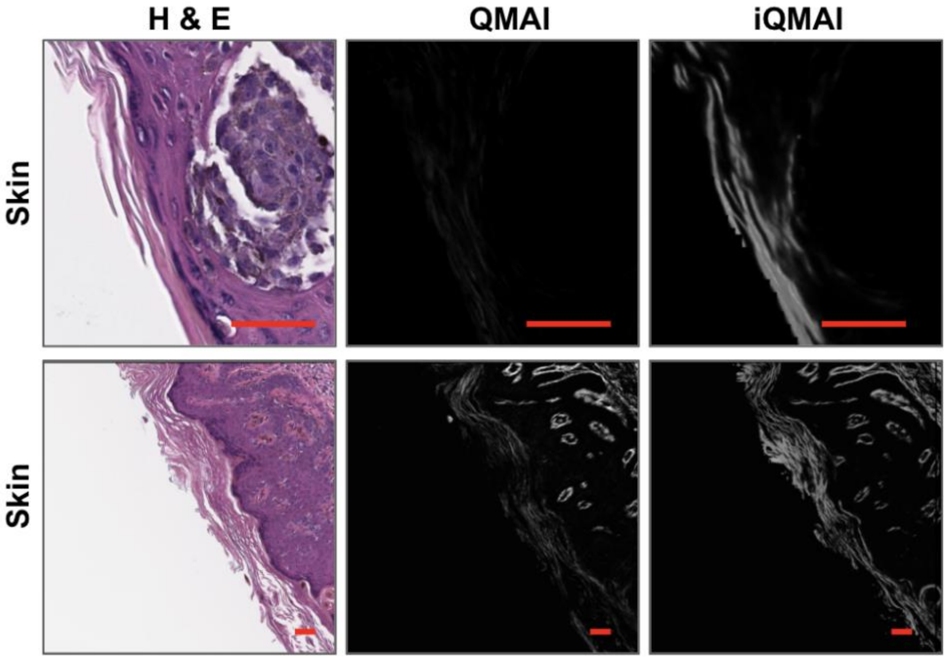


**Supplementary Figure 4. iQMAI overcall in keratin.** Scale bar, 50 μm

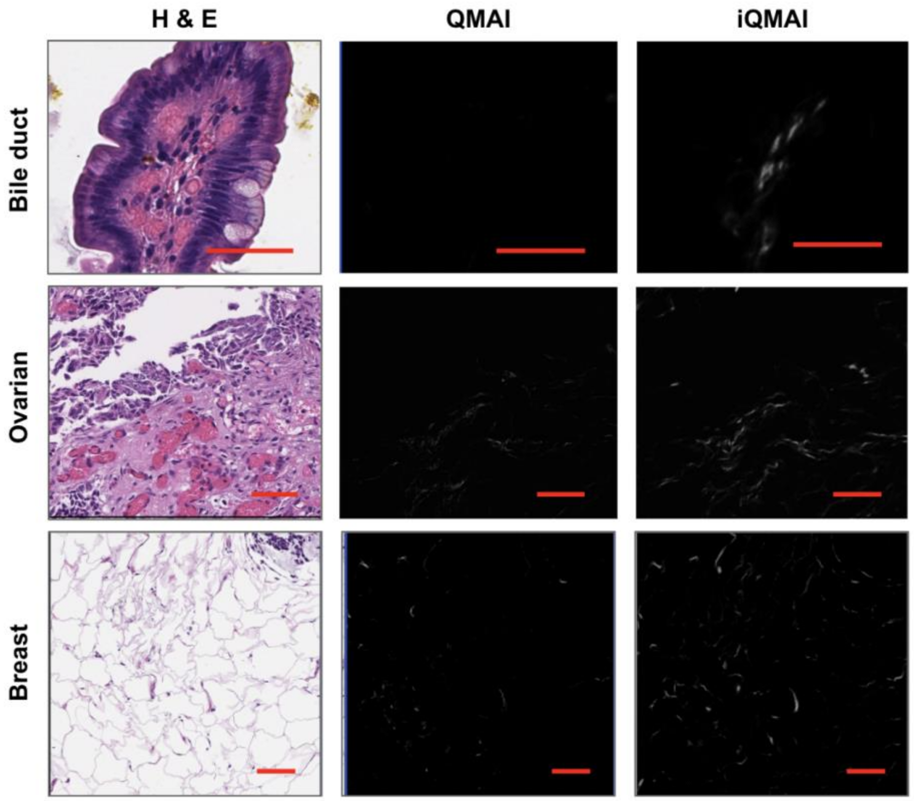


**Supplementary Figure 5**. iQMAI slightly overcalled red blood cells (top 2 rows) and walls of fat vacuoles (bottom rows). Scale bar, 50 µm

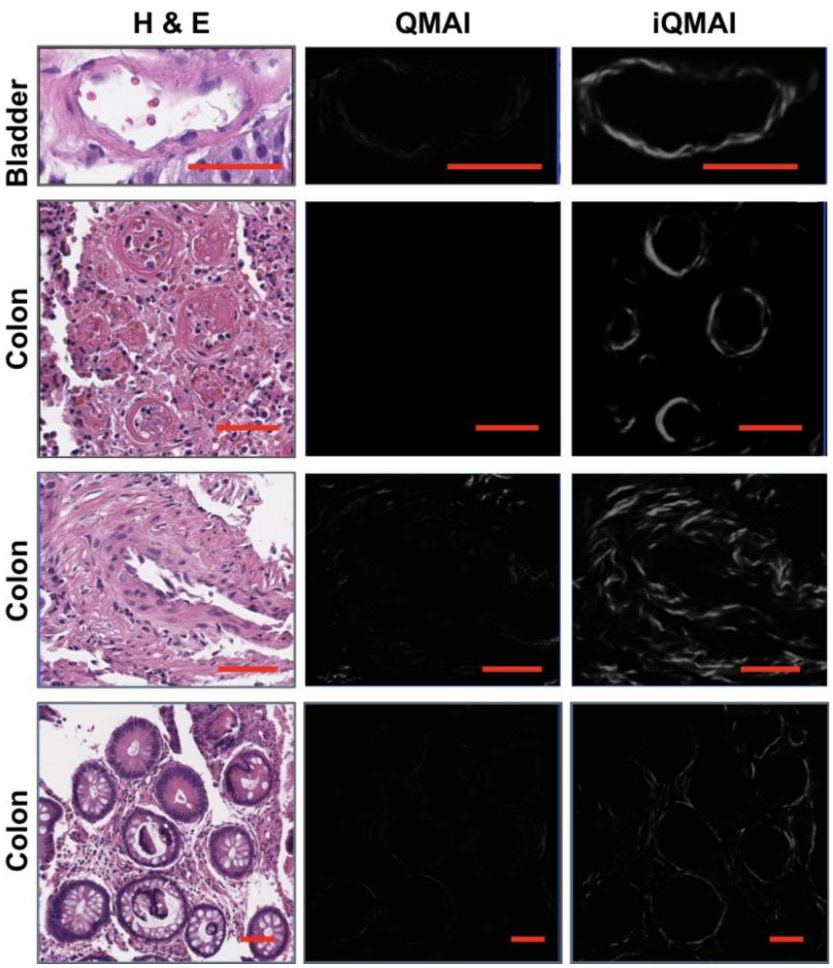


**Supplementary Figure 6.** iQMAI slightly overcalled in the wall of small vessels (top 3 rows) and basement membranes of colonic glands (bottom rows). Scale bar, 50 µm

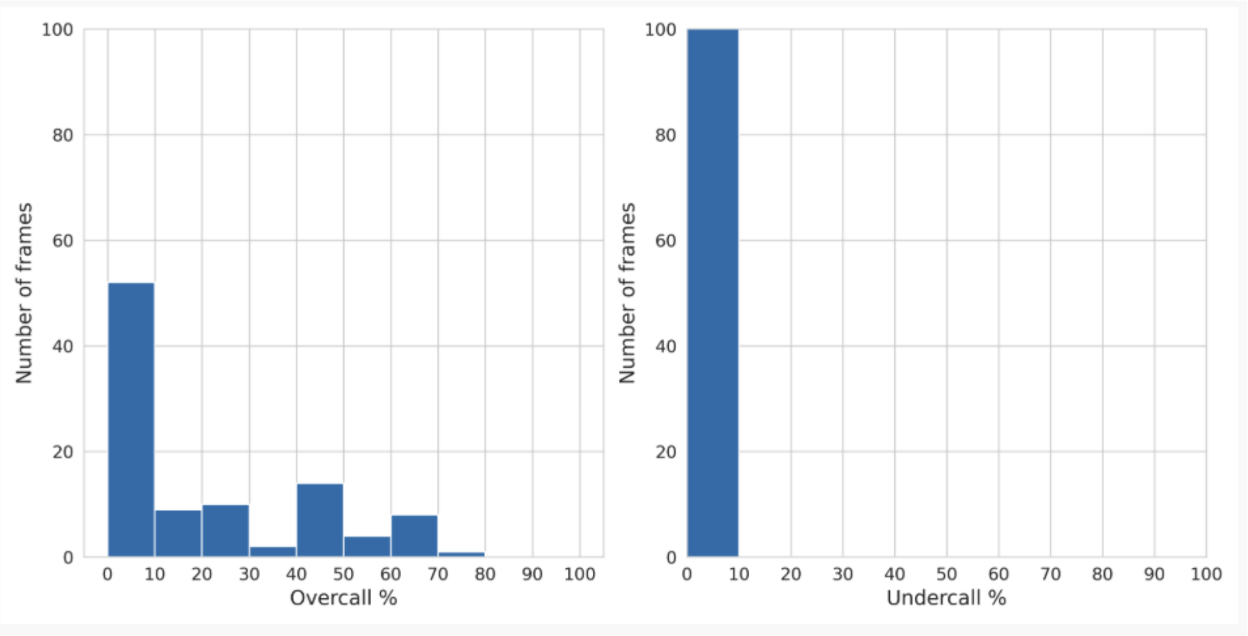


**Supplementary Figure 7.** Distribution of model overcall and undercall percentages across 100 randomly selected frames after pathologist evaluation.

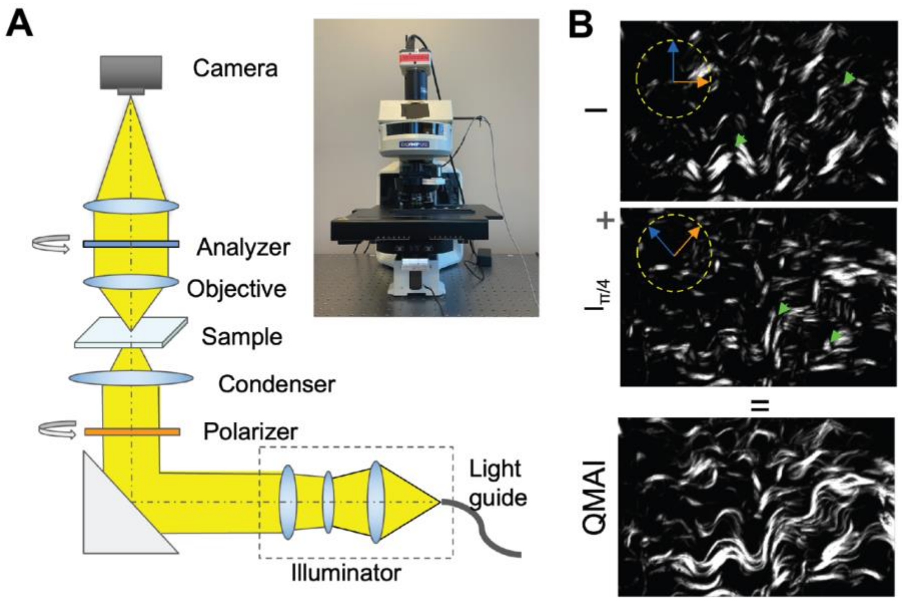


**Supplementary Figure 8. Polarization imaging to generate iQMAI training images.** A) Optical setup. The custom polarization imaging technology used herein incorporates two motorized controlled polarizers to obtain two distinct images at two angular positions (0 degrees and 45 degrees) of the transmission axis. The inset shows a picture of the real system. B) Examples of images obtained at different angular positions. The green arrowheads point to areas with reduced signals due to extinction bands. By summing these two images, the extinction band is completely removed in the image (bottom), generating a faithful representation of the collagen fibers.

### Supplementary Tables

**Supplementary Table 1.** Performance of iQMAI models ($0.7l_{LPIPS}+0.3l_{DICE}$) on the validation split with training dataset of different sizes.

| Dataset size (in millions) | ICC(2,1) | Equation | $R^2$ |
|---|---|---|---|
| 0.25 | 0.86 | y=0.56x+0.01 | 0.88 |
| 1.00 | 0.84 | y=0.54x+0.02 | 0.80 |
| 4.30 | 0.86 | y=0.56x+0.01 | 0.88 |

**Supplementary Table 2.** Performance of iQMAI models on the validation split with different reconstruction losses.

| Reconstruction loss | ICC(2,1) | Equation | $R^2$ |
|---|---|---|---|
| $l_{MAE}$ | 0.82 | y=0.57x | 0.94 |
| $l_{MSE}$ | 0.82 | y=0.57x+0.01 | 0.86 |
| $l_{LPIPS}$ | 0.83 | y=0.57x+0.01 | 0.94 |
| $0.01l_{LPIPS}+0.99l_{DICE}$ | 0.96 | y=0.83x+0.01 | 0.94 |

**Supplementary Table 3.** Performance of iQMAI models on the validation split with different weight combinations of $l_{LPIPS}$ and $l_{DICE}$

| Reconstruction loss | ICC(2,1) | Equation | $R^2$ |
|---|---|---|---|
| $1.0l_{LPIPS}$ | 0.83 | y=0.57x+0.01 | 0.94 |
| $0.7l_{LPIPS}+0.3l_{DICE}$ | 0.86 | y=0.57x+0.01 | 0.88 |
| $0.5l_{LPIPS}+0.5l_{DICE}$ | 0.92 | y=0.71x+0.01 | 0.94 |
| $0.1l_{LPIPS}+0.9l_{DICE}$ | 0.93 | y=0.72x+0.01 | 0.94 |
| $0.01l_{LPIPS}+0.99l_{DICE}$ | 0.96 | y=0.83x+0.01 | 0.94 |

**Supplementary Table 4.** Consistency between iQMAI and QMAI on fiber-based features: Slope from Linear Regression fitting

| | **Median fiber length (µm)** | | | **Median fiber relative angle (degrees)** | | | **Median fiber tortuosity** | | | **Median fiber width (µm)** | | |
|---|---|---|---|---|---|---|---|---|---|---|---|---|
| | Slope | $R^2$ | ICC(2,1) | Slope | $R^2$ | ICC(2,1) | Slope | $R^2$ | ICC(2,1) | Slope | $R^2$ | ICC(2,1) |
| Breast cancer | 1.12 | -0.21 | 0.20 | 0.96 | 0.57 | 0.78 | 0.99 | -0.08 | 0.44 | 0.89 | 0.35 | 0.73 |
| Colorectal carcinoma | 1.04 | -0.45 | 0.21 | 0.95 | 0.33 | 0.67 | 1.0 | -0.38 | 0.27 | 0.98 | 0.18 | 0.56 |
| Bladder cancer | 1.10 | -0.13 | 0.19 | 0.92 | 0.41 | 0.69 | 1.00 | -0.21 | 0.29 | 0.93 | 0.38 | 0.67 |
| Prostate cancer | 1.17 | -0.11 | 0.20 | 0.97 | 0.38 | 0.69 | 0.99 | -0.34 | 0.34 | 1.06 | 0.19 | 0.49 |
| Gastric cancer | 1.07 | -0.22 | 0.19 | 0.96 | 0.60 | 0.79 | 1.00 | -0.27 | 0.30 | 0.94 | 0.62 | 0.79 |
| Melanoma | 1.11 | -0.18 | 0.06 | 0.96 | 0.41 | 0.70 | 0.99 | -0.27 | 0.47 | 0.99 | 0.51 | 0.73 |
| Pancreatic ductal adenocarcinoma | 1.12 | -0.17 | 0.21 | 0.96 | 0.65 | 0.82 | 0.99 | -0.11 | 0.40 | 0.97 | 0.57 | 0.79 |
| Lung cancer | 1.07 | -0.19 | 0.26 | 0.94 | 0.44 | 0.70 | 0.99 | -0.18 | 0.39 | 1.00 | 0.46 | 0.72 |
| Head and neck squamous cell carcinoma | 1.05 | -0.26 | 0.17 | 0.94 | 0.57 | 0.78 | 1.01 | -0.20 | 0.39 | 1.08 | 0.38 | 0.61 |
| Cholangiocarcinoma | 1.08 | -0.38 | 0.17 | 0.96 | 0.55 | 0.77 | 0.99 | -0.12 | 0.34 | 1.00 | 0.64 | 0.80 |
| Renal cell carcinoma | 1.00 | -0.34 | 0.19 | 0.92 | 0.26 | 0.57 | 0.99 | -0.59 | 0.13 | 0.91 | 0.29 | 0.62 |
| Hepatocellular carcinoma | 1.12 | -0.17 | 0.14 | 0.94 | 0.52 | 0.75 | 0.99 | -0.77 | 0.29 | 0.97 | 0.80 | 0.89 |
| Ovarian carcinoma | 1.08 | 0.01 | 0.36 | 0.96 | 0.67 | 0.83 | 0.99 | -0.43 | 0.29 | 0.86 | 0.54 | 0.77 |
| Small cell lung cancer | 1.04 | 0.02 | 0.34 | 0.95 | 0.56 | 0.77 | 1.00 | -0.13 | 0.31 | 1.02 | 0.40 | 0.64 |
| Diffuse Large B Cell Lymphoma | 1.09 | -0.20 | 0.26 | 0.92 | 0.31 | 0.64 | 0.99 | -0.24 | 0.34 | 1.10 | 0.57 | 0.73 |

**Supplementary Table 5.** Performance of iQMAI models across slide scanners.

| | ICC(2,1) | Equation | $R^2$ |
|---|---|---|---|
| DP200 vs. AT2 | 0.92 | y=1.14x+0.01 | 0.92 |
| GT450 vs. AT2 | 0.98 | y=0.95x | 0.96 |
| C13220 vs. AT2 | 0.92 | y=1.22x+0.01 | 0.93 |

**Supplementary Table 6**. Description and quantity of samples in each slide set. Slide set A was further stratified into different splits.

| | **Set A** | **Set B** |
|---|---|---|
| Breast cancer | 46 | 3 |
| Colorectal carcinoma | 39 | 3 |
| Bladder cancer | 36 | 3 |
| Prostate cancer | 34 | 4 |
| Gastric cancer | 34 | 3 |
| Melanoma | 29 | 3 |
| Pancreatic ductal adenocarcinoma | 22 | 0 |
| Lung cancer | 21 | 1 |
| Head and neck squamous cell carcinoma | 17 | 3 |
| Cholangiocarcinoma | 15 | 2 |
| Renal cell carcinoma | 14 | 2 |
| Hepatocellular carcinoma | 13 | 2 |
| Ovarian carcinoma | 11 | 0 |
| Small cell lung cancer | 3 | 0 |
| Diffuse large B cell lymphoma | 4 | 0 |

**Supplementary Table 7.** H&E staining protocol.

| Step | Reagent | Duration |
|---|---|---|
| 1 | Xylene | 15 minutes |
| 2 | Xylene | 15 minutes |
| 3 | 100% ethanol | 2 minutes |
| 4 | 100% ethanol | 2 minutes |
| 5 | 95% ethanol | 2 minutes |
| 6 | 70% ethanol | 2 minutes |
| 7 | Water | 1 minute |
| 8 | Hematoxylin | 4 minutes |
| 9 | Water | 15 seconds |
| 10 | Water | 2 minutes |
| 11 | Define | 1 minute |
| 12 | Water | 1 minute |
| 13 | Bluing solution | 1 minute |
| 14 | Water | 1 minute |
| 15 | 95% ethanol | 15 seconds |
| 16 | Eosin | 30 seconds |
| 17 | 95% ethanol | 30 seconds |
| 18 | 100% ethanol | 30 seconds |
| 19 | 100% ethanol | 30 seconds |
| 20 | 100% ethanol | 30 seconds |
| 21 | Xylene | 30 seconds |
| 22 | Xylene | 30 seconds |
| 23 | Xylene | 30 seconds |

**Supplementary Table 8.** Tissue rehydration procedure for PSR staining.

| Step | Reagent | Duration |
|---|---|---|
| 1 | Xylene | 10 minutes |
| 2 | Xylene | 10 minutes |
| 3 | Xylene | 10 minutes |
| 4 | 100% ethanol | 2 minutes |
| 5 | 100% ethanol | 2 minutes |
| 6 | 100% ethanol | 2 minutes |
| 7 | 100% ethanol | 2 minutes |
| 8 | 95% ethanol | 2 minutes |
| 9 | 95% ethanol | 2 minutes |
| 10 | 95% ethanol | 2 minutes |
| 11 | 70% ethanol | 2 minutes |
| 12 | Reagent-grade water | 5 minutes |

**Supplementary Table 9.** Number of slides in set A that contributed to each split. For TMA slides, cores were randomly assigned to different splits.

| | Training | Validation | Test |
|---|---|---|---|
| Breast cancer | 33 | 7 | 10 |
| Colorectal carcinoma | 28 | 6 | 8 |
| Bladder cancer | 26 | 5 | 7 |
| Prostate cancer | 24 | 5 | 6 |
| Gastric cancer | 24 | 5 | 7 |
| Melanoma | 21 | 4 | 6 |
| Pancreatic ductal adenocarcinoma | 16 | 3 | 8 |
| Lung cancer | 15 | 3 | 7 |
| Head and neck squamous cell carcinoma | 13 | 2 | 4 |
| Cholangiocarcinoma | 11 | 2 | 4 |
| Renal cell carcinoma | 10 | 2 | 3 |
| Hepatocellular carcinoma | 10 | 2 | 4 |
| Ovarian carcinoma | 9 | 1 | 3 |
| Small cell lung cancer | 2 | 1 | 2 |
| Diffuse large B cell lymphoma | 2 | 1 | 4 |

**Supplementary Table 10.** Sampling results for training and validation splits.

| | Total area ($mm^2$) | Grid (μm) | Training Patches (N) | Validation Patches (N) |
|---|---|---|---|---|
| Breast cancer | 1993.0 | 80.6 | 326,971 | 21,333 |
| Colorectal carcinoma | 1032.3 | 58.0 | 316,828 | 39,412 |
| Bladder cancer | 1183.1 | 62.1 | 314,900 | 15,127 |
| Prostate cancer | 2533.4 | 90.9 | 317,062 | 10,184 |
| Gastric cancer | 510.4 | 40.8 | 310,636 | 25,011 |
| Melanoma | 612.4 | 44.7 | 301,105 | 70,226 |
| Pancreatic ductal adenocarcinoma | 2302.6 | 86.7 | 313,775 | 38,346 |
| Lung cancer | 797.6 | 51.0 | 329,189 | 7,274 |
| Head and neck squamous cell carcinoma | 808.5 | 51.3 | 337,693 | 91,437 |
| Cholangiocarcinom | 358.7 | 34.2 | 366,190 | 15,773 |
| Renal cell carcinoma | 289.0 | 30.7 | 356,269 | 24,587 |
| Hepatocellular carcinoma | 303.4 | 31.5 | 377,999 | 17,176 |
| Ovarian carcinoma | 1679.1 | 74.0 | 301,944 | 2,166 |
| Small cell lung carcinoma | 191.6 | 25.0 | 324,820 | 201,473 |
| Diffuse large B-cell lymphoma | 198.3 | 25.4 | 331,463 | 86,570 |
| **Total** | | | **4,926,844** | **666,095** |
